# The human leukemia virus HTLV-1 alters the structure and transcription of host chromatin *in cis*

**DOI:** 10.1101/277335

**Authors:** Anat Melamed, Hiroko Yaguchi, Michi Miura, Aviva Witkover, Tomas W Fitzgerald, Ewan Birney, Charles R M Bangham

## Abstract

Chromatin looping controls gene expression by regulating promoter-enhancer contacts, the spread of epigenetic modifications, and the segregation of the genome into transcriptionally active and inactive compartments. We studied the impact on the structure and expression of host chromatin by the human retrovirus HTLV-1. We show that HTLV-1 disrupts host chromatin structure by forming loops between the provirus and the host genome; certain loops depend on the critical chromatin architectural protein CTCF, which we recently showed binds to the HTLV-1 provirus. Finally, we show that the provirus causes two distinct patterns of abnormal transcription of the host genome *in cis*: bidirectional transcription in the host genome immediately flanking the provirus, and clone-specific transcription *in cis* at non-contiguous loci up to >300 kb from the integration site. We conclude that HTLV-1 causes insertional mutagenesis up to the megabase range in the host genome in >10^4^ persistently-maintained HTLV-1^+^ T-cell clones in vivo.

## Introduction

The dynamics and higher-order folding of chromatin play a critical role in gene regulation. Higher-order chromatin structure is determined by several factors, among which the best characterized is CCCTC-binding factor (CTCF) [1]. CTCF binds a non-palindromic 20-nucleotide DNA motif at ∼50,000 sites in the human genome, and acts chiefly [2] by regulating the formation of chromatin loops of ∼100 kb to ∼2 Mb, which control the contacts made between promoters and enhancers and so regulate gene expression [3–5]. Aberrant higher-order chromatin organization can result in abnormal patterns of transcription, and mutations in CTCF are linked to human disease [6–8].

Recently we found [9] that CTCF binds to a nucleotide motif in the Human T lymphotropic virus type 1 (HTLV-1; also known as the human T-cell leukemia virus) [10], when HTLV-1 is integrated - as the provirus - into the host cell genome. HTLV-1 is a primate retrovirus that infects ∼10 million people in the tropics and subtropics. The infection is asymptomatic in ∼90% of human hosts; the remaining 10% of HTLV-1-infected hosts develop either a chronic inflammatory disease, most commonly involving the central nervous system, or an aggressive malignancy of CD4^+^ T cells known as adult T-cell leukemia/lymphoma (ATL). Unlike HIV-1 infection, there remains no satisfactory treatment for either the inflammatory or malignant diseases caused by HTLV-1.

A typical host of HTLV-1 carries between 10^4^ and 10^5^ clones of infected T cells, each clone carrying a single copy of the provirus in a unique genomic location [11]. The large number of HTLV-1-infected clones appears to be established early in infection, after which persistent clonal proliferation maintains a stable hierarchy of HTLV-1-infected clones for the remainder of the host’s life. The viral regulatory proteins, Tax and HBZ (HTLV-1 bZIP), which are encoded respectively by the sense and antisense strands of the provirus, play indispensable roles in pathogenesis [12]. Tax is a transcriptional transactivator which dysregulates many host genes, while HBZ acts as a negative regulator of Tax-mediated host gene transcription and viral expression. Both Tax and HBZ contribute to persistent proliferation of the infected T cell in vivo, and it is now thought that the consequent accumulation of replicative mutations is a key driver in HTLV-1 oncogenesis. However, insertional mutagenesis has not been considered important in causing ATL.

The observation that CTCF binds to the HTLV-1 provirus [9] raised the hypothesis that CTCF bound to the provirus can form abnormal chromatin loops by dimerizing with CTCF in the flanking host genome. Using chromosome conformation capture (3C), we previously demonstrated the presence of a single CTCF-dependent chromatin loop in a long-term in vitro T cell line [9]. In the present study we extended this analysis, using circular chromosome conformation capture (4C) and RNA-seq, to examine systematically the impact of the HTLV-1 provirus on the structure and expression of the host genome in T cell clones isolated from HTLV-1-infected subjects. We show that the HTLV-1 provirus forms reproducible abnormal chromatin contacts with sites in the host genome *in cis* as far as 1.4 Mb from the provirus. Some of these abnormal chromatin contacts depend on CTCF binding to the provirus. Further, we demonstrate clone-specific deregulation of host transcription *in cis* both immediately flanking the integrated provirus (up to 50 kb upstream and downstream) and at distant sites as far as 300 kb from the provirus. Since HTLV-1 is integrated in >10^4^ different genetic locations in a typical host, and there are tens of thousands of CTCF-binding sites (CTCF-BS) in the human genome [13], these results imply that HTLV-1 has the potential to cause deregulation of host transcription at a very large number of loci in each infected host.

## Results

In this paper, we refer to loci in the host genome relative to the orientation of the HTLV-1 provirus. Thus, a locus “downstream” of the provirus is located 3′ to the 3′ LTR, whether the provirus is integrated in the positive or negative sense in the host genome.

### HTLV-1 forms chromatin loops with the flanking host genome

To test the hypothesis that HTLV-1 integration alters the host chromatin structure, we performed a genome-wide search for chromosomal positions that contact the HTLV-1 provirus, using a modified version of circular chromosome conformation capture (4C). 4C is a powerful tool to study the 3D chromatin looping between a specified genomic region (the ‘viewpoint’) with respect to the rest of the genome. The standard 4C protocol has limitations in that it is only semiquantitative [14]. To improve the quantification, we adapted our protocol for quantification of HTLV-1 integration sites [15]. We followed the 4C protocol described by van de Werken et al. [16], but instead of using a second restriction enzyme, we sonicated the library, added adapters and performed ligation-mediated PCR (see Materials and Methods). This modification confers two advantages. First, it precludes the bias towards detection of chromatin contacts that lie near a given restriction site. Second, the amplicon length serves as a unique molecular identifier, enabling relative quantification [15] of the chromatin contacts. We refer to this modified method as quantitative 4C (q4C); a similar approach (UMI-4C) has been described by others [17].

We applied q4C to test the hypothesis that the HTLV-1 provirus forms chromatin contacts with the host genome in a series of T cell clones isolated by limiting dilution from circulating CD4^+^ T lymphocytes of HTLV-1-infected individuals [18] (Table 1). Each clone has a single copy of the provirus in a unique integration site (Table S1). As the viewpoint in q4C, we used the 679 bp NlaIII fragment containing the proviral CTCF-BS (Figure 1A). Chromatin contacts were identified using a protocol based on a hidden Markov model (Materials and methods).

**Figure 1:**
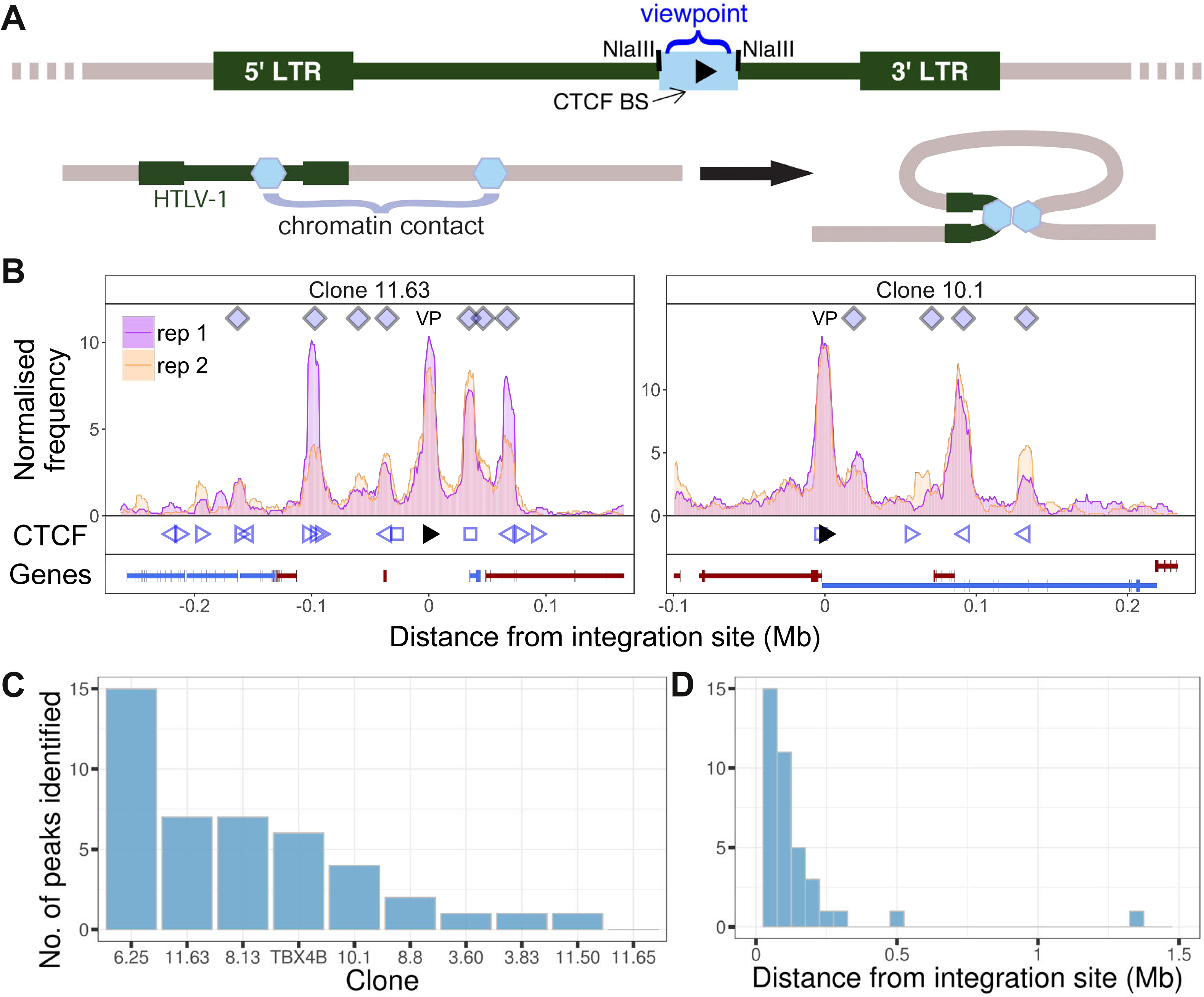
HTLV-1 forms distant contacts with the host genome. A. Upper line: the HTLV-1 genome (green), with a long terminal repeat (LTR) at each end, is integrated into a clone-specific site in the human genome (grey). The q4C viewpoint (blue rectangle) is the NlaIII fragment within the HTLV-1 genome (nucleotide residues 6564-7246) which contains the CTCF binding site (CTCF-BS; black arrowhead). Lower line: the CTCF-BS (blue hexagon) in the provirus can dimerize with a CTCF-BS in the flanking host genome. **B.** Chromatin contacts identified by q4C in 2 different clones. For each clone, the top panel depicts the q4C profile in the 5′ and 3′ host genome flanking the provirus (two biological duplicates), quantified as the normalized frequency of ligation events in overlapping windows (window width 10kb, step 1 kb). On the horizontal axis, positive values denote positions downstream of the provirus (i.e. lying 3′ of the 3′ LTR); negative values denote upstream position. VP „ viewpoint in q4C (proviral integration site). Diamonds mark the positions of reproducible chromatin contact sites called by the peak caller (Materials and Methods). CTCF panel „ open arrowheads denote positions of CTCF-BS; the filled arrowhead denotes the CTCF-BS in the provirus. Genes panel shows RefSeq protein-coding genes in the flanking host genome. The q4C profiles of remaining clones are shown in Figure S1. **C.** Number of detected peaks in each clone. **D.** Distance from detected q4C peaks to the respective proviral integration site.

We detected reproducible q4C peaks (long-range chromatin contacts between the provirus and the host genome) in 9 of the 10 infected T-cell clones examined (Figures 1B, C; Figure S1). The number of peaks per clone varied between 0 and 15, with a median of 3 peaks per clone (Figure 1C); There were significantly more peaks downstream of the integration site than upstream (p = 0.03, Chi-squared goodness-of-fit test; Figure S2A). The distance between identified peaks and the provirus of the respective clone varied between 12.9 kb and 1.4 Mb, with a median of 85 kb (Figure 1D). The distance between each peak and the integration site did not significantly differ between upstream and downstream peaks (p = 0.13, Wilcoxon test; Figure S2B).

We wished to identify whether the HTLV-1 provirus makes preferential contacts with the host genome *in cis*. The provirus is present in a single copy per cell (Figure 2A) [18]. First, to determine whether the q4C reads were derived from a single chromosome (i.e. were monoallelic) or from both homologous chromosomes (biallelic), we identified single-nucleotide polymorphisms (SNPs) present in the respective donor subject by whole-genome sequencing (see Materials and Methods). The alleles identified at heterozygous SNPs in q4C reads demonstrated that the observed ligation events were confined to a single strand (i.e. were monoallelic), with a range of at least 4 Mb from the provirus (Figure 2B).

**Figure 2:**
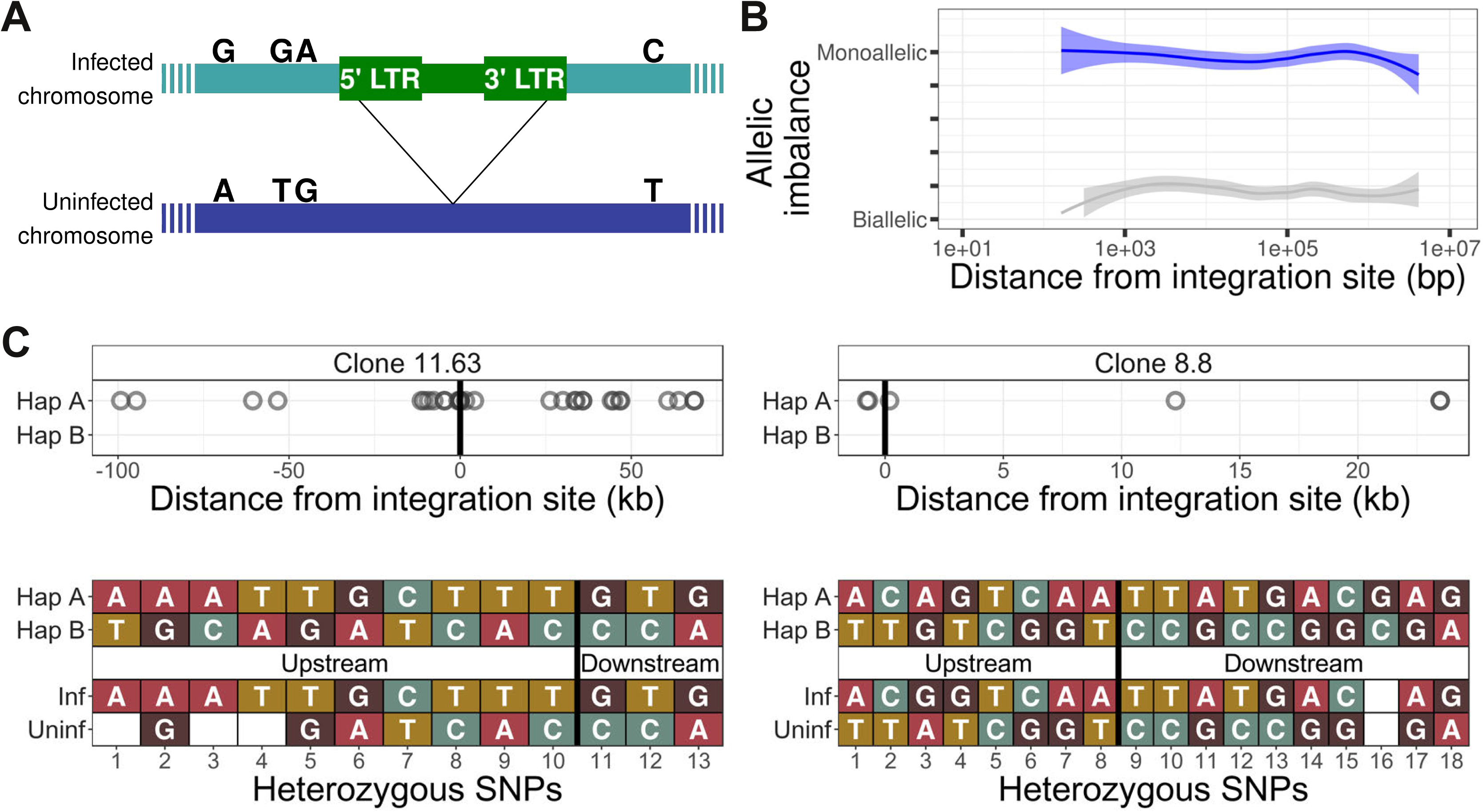
The HTLV-1 provirus makes chromatin contacts in cis with the infected chromosome. **A.** The HTLV-1 provirus is present in one copy per cell. The infected chromosome (green) can be distinguished from the uninfected homologous chromosome (dark blue) by heterozygous single-nucleotide polymorphisms (SNPs), marked by the nucleotides above each chromosome. **B.** The frequency of allele usage in unique q4C reads containing heterozygous SNPs (at least 2 reads per position) was measured, to quantify the degree of allelic imbalance, i.e. the degree of monoallelic usage present in q4C reads at a heterozygous SNP. Allelic imbalance ranges between 0 (biallelic, i.e. half of reads come from each allele) and 0.5 (monoallelic, i.e. all reads from one allele only). The dark blue line (above) shows the range of allele usage in the q4C reads; the light grey line (below) shows the allele usage for the same SNPs in the whole-genome sequencing reads. Curves were computed using LOESS regression. **C.** The infected chromosome was distinguished from the homologous uninfected chromosome using q4C data (top panel) and chromosome-specific PCR (bottom panel). Top panel - heterozygous SNPs in DNA were phased computationally to identify the two haplotypes (A and B) (Materials and Methods); the alleles present in q4C data were then assigned to the respective haplotype (circles). On the horizontal axis, positive values denote positions downstream of the provirus, and negative values denote positions upstream. Within at least 100 kb, all identified heterozygous SNP alleles mapped to only one of the two haplotypes. Bottom panel „ haplotype assignment was confirmed using haplotype-specific PCR. Each nucleotide shown is a heterozygous SNP within 5 kb of the proviral integration site, identified in the respective clone by whole-genome sequencing. The SNPs were then mapped to the respective haplotype by Sanger sequencing of long-range products amplified by PCR either between the provirus and host genome (inf „ infected haplotype) or across the provirus (uninf „ uninfected haplotype). Further examples are shown in Figure S3.

Next, we used the heterozygous SNPs to distinguish the chromosome carrying the provirus from its homologous chromosome, using computationally-determined phased haplotypes (see Materials and Methods). In this way, the infected chromosome could be distinguished from the uninfected chromosome for at least 100 kb either side of the integration site in 8 of the 10 clones (Figure 2C; Figure S3). To validate the haplotype-calling, we identified heterozygous SNPs in long-range PCR products, amplified either between the provirus and the host genome (infected haplotype) or across the proviral integration site (uninfected haplotype) (Figure 2C; Figure S3). The results (Figure 2C) showed that the reproducible contacts between the host genome and the HTLV-1 provirus were exclusively made *in cis*, that is, on the infected chromosome.

### Certain long-range chromatin contacts are CTCF-dependent

We wished to test the hypothesis that the observed abnormal long-range chromatin contacts are associated with the presence of CTCF-BS identified in the T cell clones by ChIP-seq.The results showed that that CTCF-BS were enriched at the chromatin contacts in the host genome: ∼50% of q4C peaks overlapped with at least one CTCF-BS (Figure 3A); in 10% of peaks there were two CTCF-BS. Consistent with recent findings by others [19, 20], where a peak overlapped a CTCF-BS the viral and host CTCF binding motifs were present in convergent or tandem orientation in 80% of cases (Figure 3B). The presence of a CTCF-BS was associated with a significantly greater observed q4C peak height (p = 0.025, Wilcoxon test), and there was a significant positive trend between the q4C peak height and the number of CTCF sites within the peak (Figure 3C; p = 0.016, Spearman’s test). Finally, there was a significant positive correlation between the number of observed chromatin contacts in a clone and the number of CTCF-BS within 0.5 Mb of the integration site (Figure 3D; Pearson’s correlation test); this correlation remained significant up to 1.16 Mb from the provirus.

**Figure 3:**
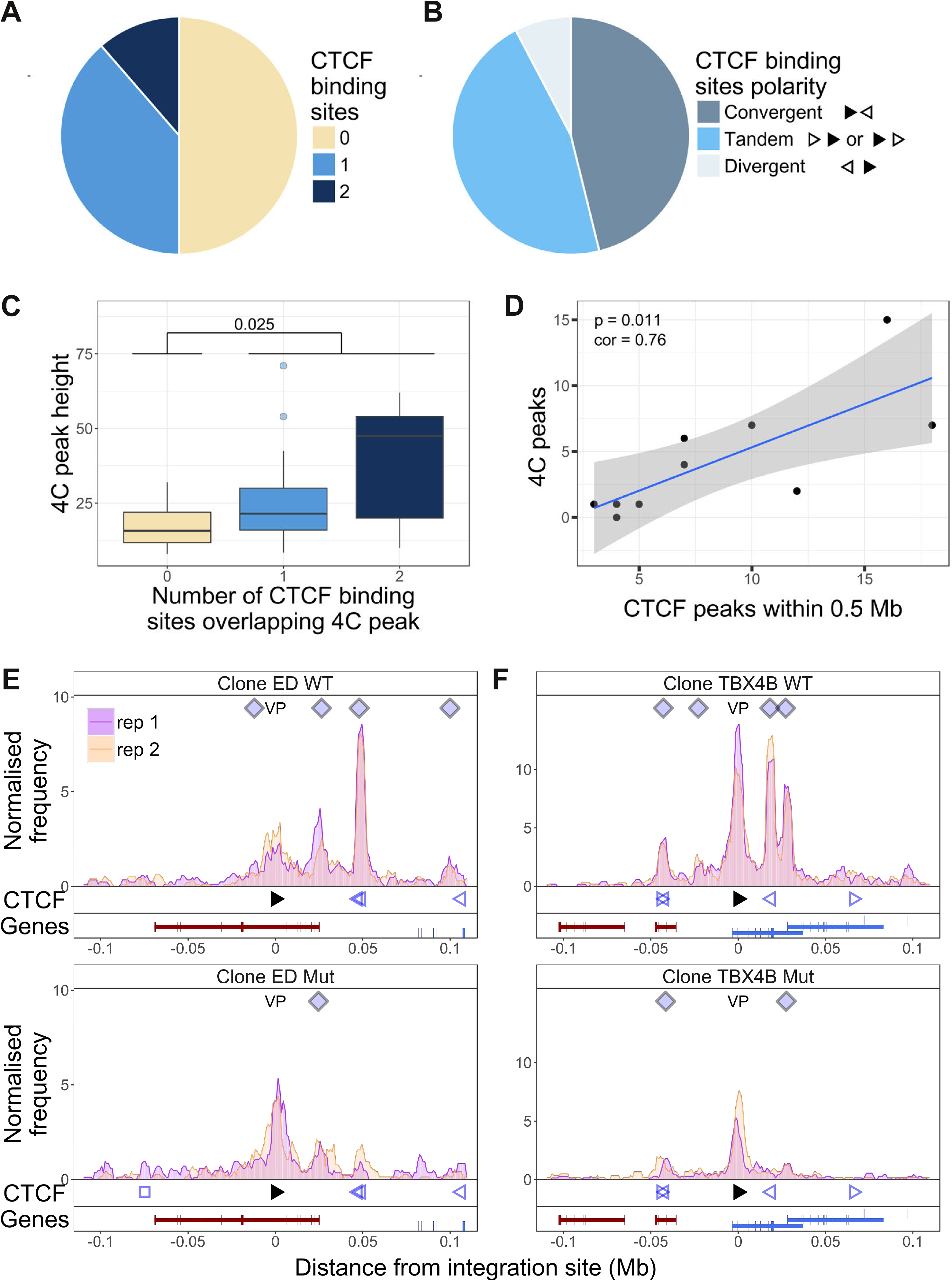
Dependency of virus-host contacts on CTCF binding. **A.** Of 44 contacts identified by q4C in the clones examined, 22 contained one CTCF-BS (N = 17) or two CTCF-BS (N = 5); the remaining 22 contacts did not contain a CTCF-BS. **B.** The polarity was determined of the proviral CTCF-BS (filled arrowhead) and the host CTCF-BS (open arrowhead). Of the CTCF-containing peaks whose polarity could be determined, convergent orientation (possible only for downstream peaks) was found in 46% of peaks, divergent orientation (possible only for upstream peaks) in 8% of peaks, and tandem orientation (possible for either upstream or downstream peaks) in 46% of peaks. **C.** Distribution of q4C peak height (mean number of ligation events between replicates identified in each peak) in peaks containing 0, 1 or 2 CTCF-BS (coloured as in panel A): peaks that contained at least one CTCF-BS were significantly higher than those that lacked a CTCF-BS (p = 0.025, Wilcoxon test). In addition, there was a significant correlation between mean q4C peak height and the number of overlapping CTCF-BS (p = 0.0156, Spearman’s rank correlation test) (not illustrated). **D.** The number of observed contacts was positively correlated with the number of CTCF-BS within 0.5 Mb of the proviral integration site (p = 0.011, Pearson’s correlation test). **E-F.** q4C analysis was carried out on a clone from an ATL-derived cell line (E) and a T-cell clone (F), respectively either the wild-type cells (WT; top panels) or after CRISPR-Cas9 knockout of the proviral CTCF-BS (bottom panels). The vertical axis shows the normalized number of q4C ligation events (overlapping windows 5 kb wide, with 1 kb steps). The bottom track in each panel shows the position of known RefSeq protein-coding genes; in clone TBX4B the provirus is inserted between exons of the gene *PNPLA* (shown in blue; see Results section).

We then tested whether CTCF binding to the provirus is required for formation of provirus-host contacts. We used a CRISPR-modified cell line (ED) [9], derived from an individual with the HTLV-1-associated malignancy adult T-cell leukemia (ATL) (Figure 3E). We also applied CRISPR-Cas9 ribonucleoprotein transfection [21] to knock out the CTCF-BS in a non-malignant T-cell clone (TBX4B; Figure 3F). We carried out q4C analysis on cells from the wild-type (WT) clone, and the mutant (Mut) clone containing a mutated CTCF-BS. The results show that the loss of CTCF binding was associated with a loss of 5 of the 8 observed contacts; three of the 5 lost contacts overlapped a CTCF-BS in the host.

### HTLV-1 alters contiguous host transcription

The results obtained by q4C demonstrated that HTLV-1 integration can alter host chromatin looping. We wished to investigate the impact of the HTLV-1 provirus on transcription in the host genome both immediately flanking the provirus and at distant loci. We carried out strand-specific mRNA-sequencing (RNA-seq) on the HTLV-1-infected T cell clones, and quantified the density of reads mapping to discrete 1 kb windows up to 2 Mb from the respective proviral integration site. The results showed upregulated transcription in the host genome immediately flanking the HTLV-1 provirus, in all clones examined (Figure 4A, B; Figure S1, S4). This upregulated transcription was observed either upstream or downstream of the provirus, or both, with a predominant increase downstream in the same sense as the HTLV-1 plus-strand and upstream in the opposite sense (Figure 4C). We then performed this analysis separately to compare the clones with high HTLV-1 plus-strand expression and those with low plus-strand expression (defined respectively as those clones with a *tax* read intensity in the RNA-seq above or below the median intensity of all clones). The results showed that, whereas abnormal upstream antisense expression was present in most clones, an increase in same-sense transcription was specific to those clones with high plus-strand expression (Figure 4D).

**Figure 4:**
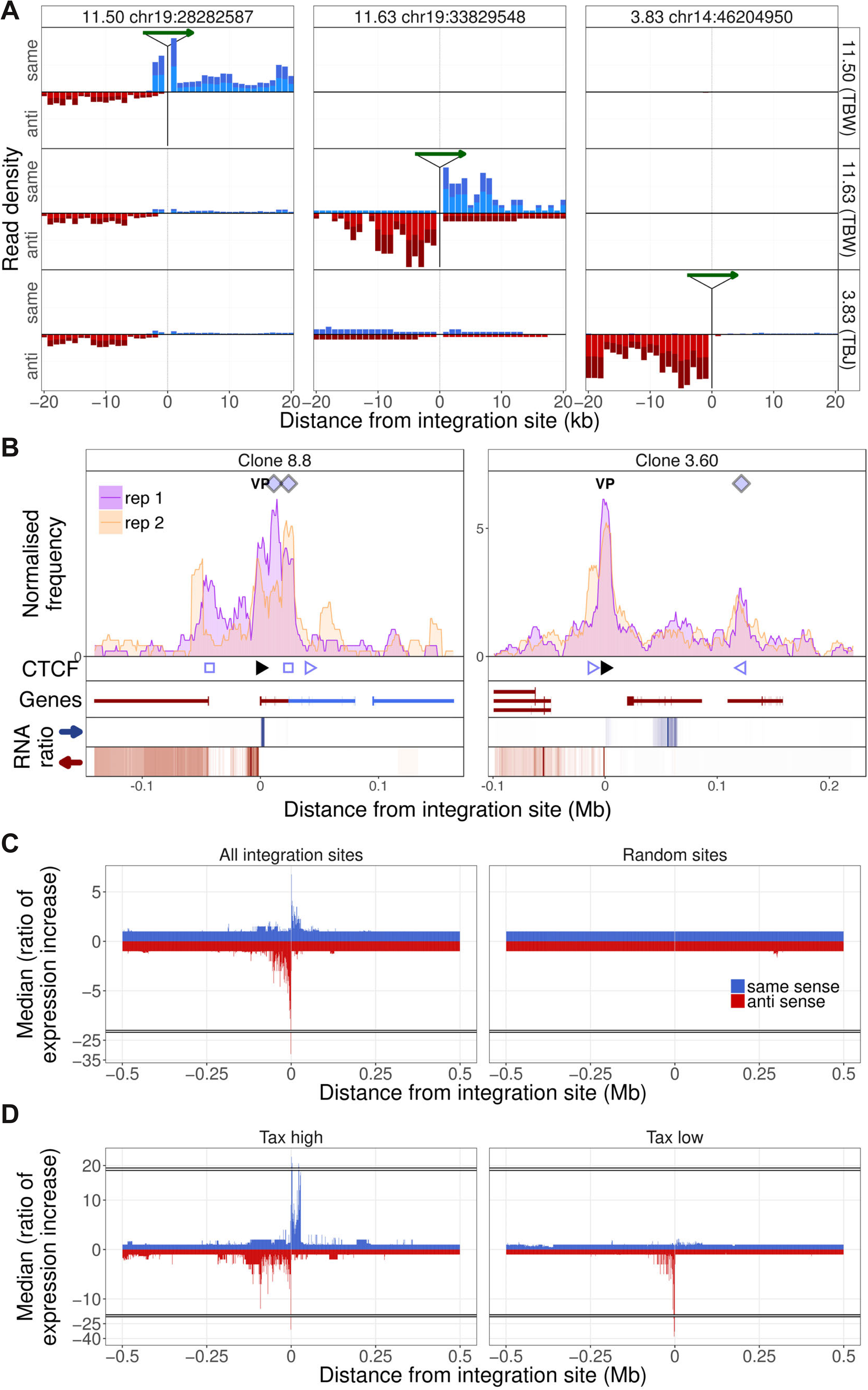
Integration site-specific upregulation of host transcription. **A.** In each column, the green arrow indicates the HTLV-1 proviral integration site in the clone indicated at the top of the column. Each row shows the the transcription density (normalized RNA-seq read count) flanking that genomic position in the clone indicated at the right-hand side. In each case, transcription orientation and positions are shown relative to the integrated provirus. Read density shown in blue shows transcription in the same orientation as the proviral plus-strand; red shows transcription in the antisense orientation to the proviral plus-strand. **B.** q4C profiles of two clones aligned with the transcription density within 300 kb of the proviral integration site. The RNA ratio shows the ratio of transcription density in a given bin (number of reads in 1 kb bin / total number of reads in sample) in the target clone, divided by the median expression density of all clones in that bin. Colours represent expression in the same sense (blue) or opposite sense (red) to the HTLV-1 plus-strand. Data on the remaining clones are shown in Figure S1. **C.** Left panel: median ratio of transcription density of all clones, aligned on the integration site (1 kb bins, up to 0.5 Mb from the integration site). Right panel: median ratio of transcription density at 10 genomic positions, selected at random from a gap-excluded hg19 reference genome. **D.** Analysis carried out as in panel C, separately for clones expressing HTLV-1 plus-strand transcripts at a level greater than (left panel) or less than (right panel) the median of all clones.

The observed upregulation of transcription was frequently intergenic, but we also observed instances of clone-specific gene expression. For example, in clone TBX4B, HTLV-1 is inserted between exons of the gene *PNPLA* (Figure 3F). This gene was not expressed in the other T cell clones, but was highly expressed in TBX4B both downstream and upstream of the integration site, in the same sense as the proviral plus-strand. The presence of abnormal same-sense host transcription upstream of the 5′ LTR suggests that the transcription was driven by the proviral enhancer.

### Transcription is altered at non-contiguous sites

In addition to the abnormal transcription immediately flanking the provirus, there were frequent examples of clone-specific transcription in regions of the host genome not contiguous with the provirus (Figure 4B. Figure S4). For example, in clone 8.8, there was upregulation of host transcription both flanking the integration site and in the gene *TBC1D4* (a Rab GTPase-activating protein) ∼44 kb upstream; no transcripts were detected in the intervening host genome (Figure 4B, left). Non-contiguous transcription also occurred at the downstream contact site made with the provirus, resulting in aberrant splicing of the gene *UCHL3* to produce putative novel *UCHL3* transcripts (Figure S5B).

In clone 3.60, abnormal non-contiguous transcription was found downstream of the provirus (Figure 4B, right) between the provirus and the contact (indicated by the q4C peak) formed with the host genome. This region contains a gene (*SULT1B1*) in the negative strand of the genome; the abnormal transcription observed in clone 3.60 was present in the positive strand with alternative splicing, resulting in a putative novel transcript (Figure S5A). These observations demonstrate that the provirus can alter transcription *in cis* both within and between genes, and alter splicing, producing novel transcripts.

### HTLV-1 alters host transcription *in cis*

To test the hypothesis that the observed abnormal transcription of the host genome was confined to the chromosome carrying the provirus, we quantified the allelic imbalance in the SNPs identified in RNA-seq reads up to 2 Mb from the integration sites. The results showed that in heterozygous SNPs identified in the clone-specific transcripts, transcription was predominantly homozygous, i.e. monoallelic (Figure 5A).

**Figure 5:**
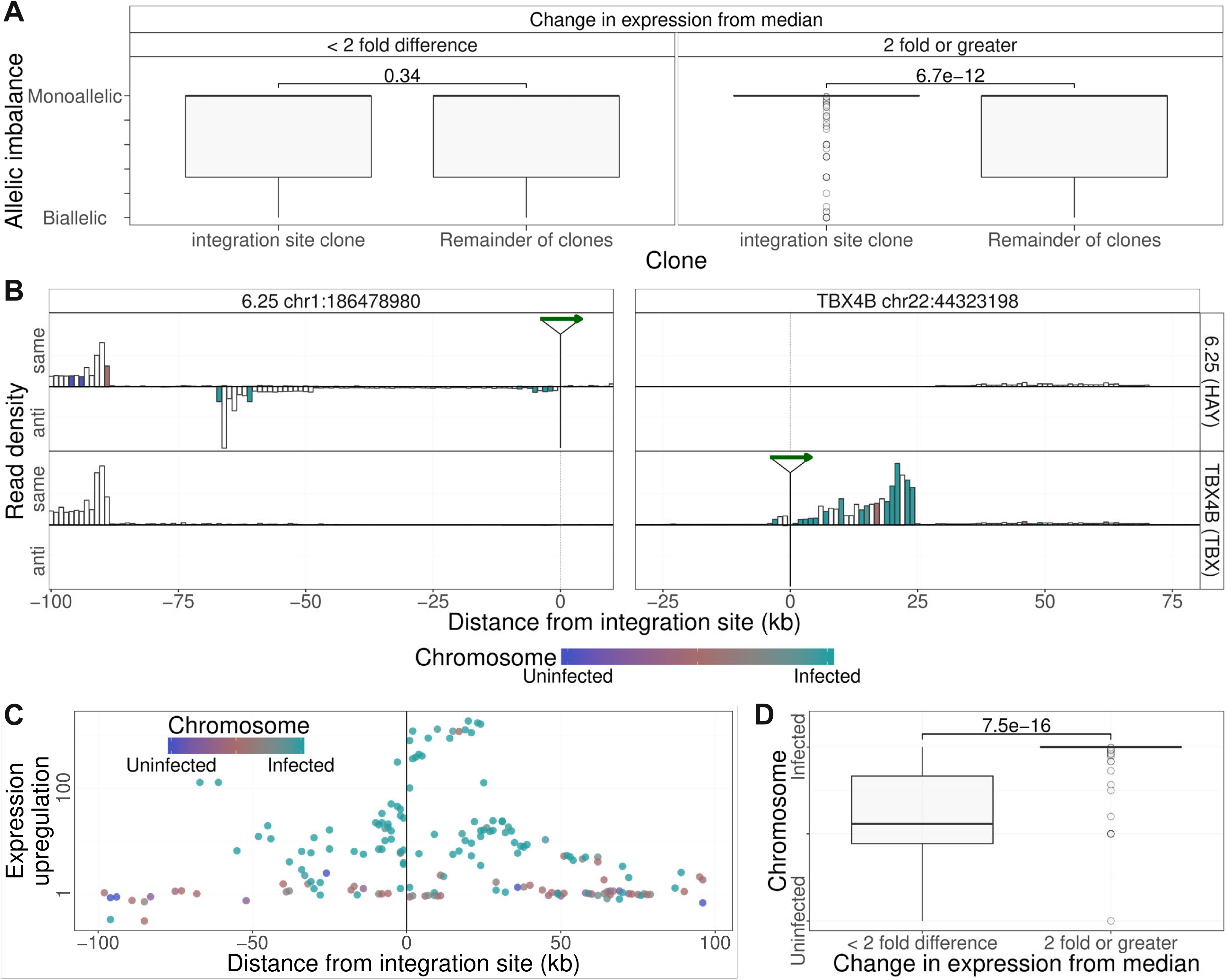
Clone-specific host transcription is derived from the infected chromosome. **A.** Allelic imbalance (AI) denotes the degree of monoallelic usage of identified SNPs: AI =0 indicates biallelic transcription; AI = 0.5 indicates monoallelic transcription. In each clone, the AI was quantified in transcripts within 2 Mb of the proviral integration sites and compared with the value at that site in all other clones. Clone-specific transcription (transcription density in the clone carrying the provirus, 2-fold or greater than the median; 1 kb bins) was monoallelic; shared transcription was biallelic. While there was no significant difference between the allelic imbalance in those bins for which there was little or no change in transcription from median, for those bins where clone specific expression was observed (2 fold or greater increase), the allelic imbalance was significantly greater (more monoallelic) in the integration site clone compared to remainder of clones (p = 6.7 * 10^−12^, Wilcoxon test). **B.** Transcription density depicted as in Figure 4A, analysed by haplotype (see Figure 2). Columns are coloured by the mean frequency of infected or uninfected alleles (1 kb bins). White columns did not include SNPs that could be assigned to a single haplotype. **C.** Median ratio of transcription density (log scale) in 1 kb bins containing a heterozygous SNP coloured by the frequency of alleles derived from the infected (green) or uninfected (blue) haplotypes. **D.** The SNP alleles expressed at = 2 × median level were over-represented in the infected haplotype.

To test whether this monoallelic transcription was derived from the infected chromosome or its homologous chromosome, we used the haplotype-calling approach described above to assign heterozygous SNPs present in the RNA-seq reads within 100 kb of the proviral integration site to either the infected or the uninfected chromosome. The results (Figure 5B, C, D), showed that, where the transcripts could be assigned to one chromosome, the observed clone-specific transcripts were overwhelmingly derived from the infected chromosome, whereas transcripts that were not specifically upregulated were expressed from either chromosome at a similar frequency (Figure 5B, left side of left panel; Figure 5C, D).

**Table 1:** T cell clones used.

| subject | clone(s) |
| --- | --- |
| TBJ | 3.60, 3.83 |
| TCX | 8.13, 8.8 |
| TCT | 10.1 |
| TBW | 11.50, 11.63, 11.65, 13.50(U) |
| TBX | TBX4B |
| HAY | 6.25, 6.30(U) |

## Discussion

The mammalian genome is not randomly arranged in the nucleus, but is folded in a highly ordered manner at a series of successively larger spatial scales [22]. At the smallest scale, chromatin is organized in a series of reproducible loops that are formed by bringing together specific genomic loci which are separated on the linear genome by up to ∼2 Mb [23–25]. The zinc finger protein CTCF plays a central part in establishing and maintaining these chromatin loops: the non-palindromic CTCF-BS is found at the borders of ∼80% of loops. Certain other chromatin-associated proteins, such as PRC1 [26], and transcription factors including AP-1 [27] and YY1 [28], can also cause looping of chromatin. The resulting chromatin loops in turn play a critical part in controlling gene expression, by regulating the contacts made between enhancers and promoters [3–5]. Disruption of chromatin loops can deregulate gene expression and cause developmental abnormalities [29] or diseases such as cancer [6–8].

The discovery that the HTLV-1 provirus binds CTCF [9] therefore raised the hypothesis that the provirus forms chromatin loops with the neighbouring host genome and thereby deregulates host transcription. To test this hypothesis, we used a panel of non-malignant CD4^+^ T cell clones naturally infected with HTLV-1. Each clone carries a single copy HTLV-1 provirus in a known genomic location [18]; all 10 clones studied were competent to express the plus-strand proviral genes except clones 6.25 and 8.13.

The results of chromosome conformation analysis (q4C) presented here reveals reproducible contacts between the HTLV-1 provirus and the flanking host genome at least 1.3 Mb from the integration site. Each clone has a unique pattern of chromatin contacts, depending on the site of integration of the provirus in the host genome. The allelic bias observed in the q4C data, with the preferential detection of SNPs on the chromosome carrying the provirus, indicates that the provirus makes preferential contacts *in cis* with host chromatin at distances at least 4 Mb from the provirus. Preferential contacts *in cis* may extend beyond this distance, but detection is likely to be limited by the sensitivity of the q4C technique.

Four observations indicate that certain chromatin contacts made between the provirus and the host genome depend on the presence of CTCF. First, in each of two clones (ED and TBX4B; Figure 3E, F), knocking out the CTCF BS in the provirus without altering the coding sequence of the *tax* gene abrogated a major contact with a site in the host genome, respectively ∼50 kb and 17 kb downstream, where CTCF was shown to bind. Second, of the 44 host loci identified by the peak-calling algorithm as making contact with the provirus, 22 contained one or more CTCF-BS. Third, consistent with recent observations made by others [19, 20], 12 of the 13 (92.3%) CTCF-BS in the host contact sites whose orientation could be ascertained were oriented towards the CTCF-BS in HTLV-1. Fourth, both the number and height of q4C peaks were correlated with the number of CTCF-BS in the host genome (Figure 4C, D). However, not all host chromatin contact sites contain CTCF-BS, and certain peaks remained detectable by q4C after knock-out of the proviral CTCF-BS. These observations are consistent with the findings that certain chromatin-associated proteins other than CTCF can also give rise to chromatin looping [26–28].

The formation of chromatin loops between the HTLV-1 provirus and the host genome in turn raised the possibility that host transcription is deregulated. At least two mechanisms of such deregulation can be suggested. First, the abnormal chromatin loops might alter the normal contacts made between enhancers and promoters in the host genome, either creating new contacts or abrogating pre-existing contacts. Second, the new chromatin loop might bring the proviral LTR near a host promoter and cause abnormal transcription from the promoter.

We show here that HTLV-1 can deregulate host transcription both at sites contiguous with the provirus and at non-contiguous, distant sites. Again, each clone has a unique pattern of transcription. In the host genome immediately flanking the provirus, we detected frequent abnormal (clone-specific) transcription, predominantly in the opposite transcriptional sense to the provirus upstream of the 5′ LTR and, to a lesser extent, in the same transcriptional sense downstream of the 3′ LTR. The observed abnormal transcription involved both intergenic regions and identified host genes (Figure 4B, Figure S1, S4); in addition, we observed examples of abnormal transcription and abnormal splicing of host genes (Figure S4).

Kataoka et al. [30] and Rosewick et al. [31] have reported evidence of transcription initiated on the minus strand of the provirus and continuing upstream into the host genome in transformed lymphocyte clones carrying HTLV-1 or the related bovine leukemia virus (BLV). The present results demonstrate that abnormal host transcription flanking the provirus *in cis* is a general feature of HTLV-1 infection. This abnormal transcription is bidirectional: both downstream of the provirus in the same sense as the proviral plus-strand, and upstream in the opposite sense. The observation of clone-specific transcription upstream of the 5′ LTR in the plus-strand sense (Figure 4A), especially in high *tax*-expressing cells (Figure 4D), suggest that the HTLV-1 promoter/enhancer can drive abnormal expression from nearby transcription start sites.

In addition to the abnormal transcription frequently detected in regions of the host genome contiguous with the provirus, each clone also had a unique pattern of abnormal transcription at non-contiguous, distant sites. The great majority of these clone-specific transcripts came from the chromosome carrying the provirus, not its homologous “uninfected” chromosome, as shown by the allelic bias observed in the sequence reads (Figure 5D). Similarly, the observed allelic bias in the q4C reads extended to at least 1 Mb upstream and downstream of the provirus (Figure 2B). These results are consistent with the notion that the abnormal transcription observed at sites distant from the provirus results from contacts made between the provirus and the host genome.

While certain CTCF-dependent chromatin loops mediate enhancer-promoter interactions, there is growing evidence that the main function of CTCF-dependent loops is structural and is largely shared between cell types, whereas loops formed by other chromatin-binding proteins such as YY1 [32] or chromatin-modifying proteins such as PRC1 [33] mediate dynamic enhancer-promoter interactions and thus play a central part in determining cell-type-specific transcription. It remains to be tested whether each respective instance of HTLV-1-associated abnormal host transcription observed here depends on chromatin looping induced by CTCF or another chromatin-binding protein.

Retroviruses have long been known to disrupt host gene expression by insertional mutagenesis, either by integration of the provirus within a gene or by activating expression of a flanking host gene [34–36]. Upstream host oncogenes have been shown to be activated by integrated retroviruses, for example mouse mammary tumour virus [36] and murine leukemia virus; in the latter case transcription was initiated in the LTR and read through into the host genome [37]. Integration downstream of a host oncogene conferred a strong selective advantage on certain clones transduced with a gene therapy vector derived from MLV, resulting in a number of cases of leukemia [38, 39]. In mice, it has also been shown that MLV can cause abnormal host transcription by activating a distant host gene [40–42]. In a study of MLV-induced leukemia, Sokol et al. [41] suggested that the MLV proviral enhancer was brought near the oncogene by chromatin looping; if so, it is likely that chance integration of MLV at this particular locus enabled the virus to exploit a normal chromatin loop normally present in the mouse genome. In contrast to this adventitious effect of MLV, our results show that the HTLV-1 provirus itself can cause both abnormal chromatin contacts *in cis* with distant host loci and abnormal host transcription at distant loci. These findings demonstrate that HTLV-1 has a range of insertional mutagenesis that can extend to the megabase scale.

Schmidt et al [43] showed that retrotransposons propagated CTCF-BS throughout the genome of several mammalian lineages during evolution; however, present-day exogenous retroviruses have not been shown to alter the higher-order structure of host chromatin, to our knowledge. It was recently reported [44] that the spumaretrovirus foamy virus encodes a CTCF-BS in its long terminal repeats. The impact of these binding sites on host chromatin structure and transcription have not been investigated; Goodman et al [44] suggested that the presence of CTCF bound to the LTRs may block enhancer effects of the foamy virus LTR, and so account for the low observed degree of foamy virus genotoxicity.

The HTLV-1 transcriptional transactivator protein Tax drives expression not only of the proviral plus stand but also of several host genes, notably *CD25* (*IL2RA*), *IFNG, IL6, IL15, GM-CSF, TNFB*, and *CCL22* [45]. While these effects may contribute to the persistence and replication of the virus, and to the risk of the inflammatory and malignant diseases associated with HTLV-1, the effects are not clone-specific. The present results show that, in addition to these generic effects, HTLV-1 has the potential to disrupt host gene expression *in cis* at both contiguous and non-contiguous sites. Since the virus infects between 10^4^ and 10^5^ different clones of T cells in a typical host, each carrying the provirus at a different genomic location, we conclude that the virus deregulates tens of thousands of host genes in a typical infected host.

## Materials and Methods

### Cell culture, preparation

The HTLV-1-infected T-lymphocyte clones were derived by limiting dilution from peripheral blood mononuclear cells (PBMCs) of donors attending the National Centre for Human Retrovirology (NCHR) at Imperial College Healthcare NHS Trust, St Mary’s Hospital, London. All donors gave written informed consent in accordance with the Declaration of Helsinki to donate blood samples to the Communicable Diseases Research Tissue Bank, approved by the UK National Research Ethics Service (15/SC/0089). The derivation of these clones and the genomic insertion site of the single-copy HTLV-1 provirus in each clone were previously reported [18]. The cells were cultured in RPMI-1640 medium (Sigma-Aldrich) with added L-glutamine (Invitrogen), penicillin and streptomycin (Invitrogen), 10% AB human serum (Invitrogen) at 37°C, 5% CO_2_. IL-2 (Promokine) was added to the culture every 3 days, and the retroviral integrase inhibitor raltegravir (Selleck) was maintained at 10 µM throughout cell culture, to prevent secondary infection. In addition, the cells were activated every 14 days by the addition of beads coated with antibodies against CD2, CD3 and CD28 (Miltenyi-Biotech). All experiments were carried out on cells harvested on Day 9 of this cycle, after addition of fresh media on Day 8.

### CRISPR/Cas9 modification

We used Cas9 ribonucleoprotein transfection [21] to selectively mutate 6 nucleotides in the core CTCF binding-site in the HTLV-1 provirus and abrogate CTCF binding [9].

### q4C

q4C„seq libraries were prepared according the 4C protocol by [16] and our protocol for linker-mediated PCR (LM-PCR) [46] with a modification. Eight million cells were crosslinked in phosphate-buffered saline (PBS) containing 1% formaldehyde for 10 min at room temperature. Cells were lysed using a buffer containing 10 mM Tris-HCl (pH 8.0), 10 mM NaCl, 0.2% NP-40 and complete EDTA-free Protease Inhibitor Cocktail (Roche). The DNA was digested with NlaIII (New England Biolabs, NEB) in the presence of 0.2 % SDS, followed by ligation using T4 DNA ligase (NEB) under dilute conditions. The ligated DNA (3 µg) was sheared by sonication with a Covaris S2 instrument; adapters were added and the fragments were subjected to LM-PCR for preparation of 4C–seq libraries. DNA fragments were end-repaired using T4 DNA polymerase, DNA polymerase I Klenow fragment, and T4 polynucleotide kinase (NEB). An adenosine residue was added at the 3′ end of the DNA fragments using Klenow fragment 3′ to 5′ exo– (NEB). A partially double-stranded DNA linker with a specific 6-bp tag was ligated to the DNA ends using a Quick ligation kit (NEB). The linker-ligated product (200 ng per reaction tube) was amplified by a first PCR (PCR1) using the primers HY3 (5′-CTCCTCCTTGTCCTTTAACTCTTCCTC-3′) and Bio4 (5′-TCATGATCAATGGGACGATCA -3′) and Phusion DNA polymerase (NEB). Eight individual PCR1 reactions of 50 µL were prepared for each sample and purified individually using QIAquick PCR Purification Kit (Qiagen) columns. To perform PCR2, 1/150th of the purified PCR1 product was amplified between the primers HY12 (5′-AATGATACGGCGACCACCGAGATCTACACCCTCCAAGGATAATAGCCCGTC-3′) and P7 (Illumina). The 8 PCR2 products were combined and purified by QIAquick PCR Purification Kit (Qiagen). The libraries were quantified by qPCR using Illumina primers P5 and P7. Stock 4C libraries were diluted accordingly and clustered on the sequencing flow cell.

The steps in q4C analysis are summarized in Figure S6. q4C libraries were sequenced with paired ends (read1 and read 2), with a read length of 100 bp with 8 initial dark cycles plus a 6-bp tag read (read 3), on a HiSeq 2500 in Rapid-run mode (Illumina). The sequencing primer was situated in the viewpoint (NlaIII fragment), terminating 4 bases upstream of the 3′ NlaIII restriction site. These 4 bases as well as the NlaIII restriction site were used as a filter to ensure sequence specificity, and were subsequently trimmed from the 5′ end of the sequence in the first read of each pair using cutadapt [47]. Adapter and low quality bases were trimmed using trim galore (http://www.bioinformatics.babraham.ac.uk/projects/trim_galore/). In order to avoid misalignments due to composite reads (containing NlaIII fragments from multiple contacts), in silico digestion at NlaIII binding sites (CATG) was performed on the reads. ‘Digested’ Fastq files were subsequently aligned (as single reads) against a merged reference of the human genome (hg19 assembly) and the HTLV-1 genome (accession number AB513134). For each read, only the first digested NlaIII fragment was used for subsequent analysis unless the first fragment was the one directly following the viewpoint (incomplete digestion) in which case the second fragment was used. Successfully aligned pairs were filtered to remove barcode errors, and only pairs where both read 1 and read 2 mapped to the same chromosome in convergent orientation were kept. Unique read1-read2 pairs were considered a ligation event: the shear site serves as a unique molecular identifier [46], i.e. it identifies a ligation event in a single cell. The number of unique ligation events was then quantified at each ligation site (virus-host genome contact site). At least two biological replicates were analysed from each clone. Where more than 2 replicates were analysed, the two with the highest library diversity (the highest total number of ligation events) were used in subsequent analyses.

To call chromatin contact sites from the q4C data, we first counted the unique ligation events (comprising distinct ligation and shear positions) in 5 kb overlapping windows (1 kb steps) across the alignments for every clone from each sample. Next we trained a three-state hidden Markov model on all windows with more than one ligation event, for chromosomes containing integration sites, ordered by genomic position, using the Expectation-Maximization (EM) algorithm from the depmixS4 package [48] to find the initial parameters. We then applied the Viterbi algorithm using this trained model to all individual clones and samples separately.

To define continuous genomic regions of interactions for a given chromosome, we applied a cubic smoothing spline to each sample’s respective state space, interpolating over n points across the chromosome with n being 0.25 * the number of overlapping windows in that chromosome. The edges for each peak were defined by the change in sign of the difference of the curve between consecutive points. Peaks were called for a single replicate only if included windows with a minimum of state 2 (probable peak) in both replicates. The peaks were intersected between the replicates using the GenomicRanges package [49] and intersects which did not include the local maximum of each peak in both replicates were discarded. Peaks which were unique to one clone, were under 50 kb in width and did not overlap the integration site were used in subsequent analysis.

### RNA-seq

RNA-seq libraries were prepared from total RNA of the T cell clones using the TruSeq Stranded mRNA HT Sample Prep Kit according to the manufacturer’s instructions, and sequenced with the HiSeq4000 (150 bp paired-end reads). RNA sequencing for each clone was performed using the Ilumina platform. Where more than one replicate was sequenced, the one with a larger number of reads was used in subsequent analysis. Data quality was assessed using FastQC (https://www.bioinformatics.babraham.ac.uk/projects/fastqc/) and aligned to the same combined reference (human genome + proviral genome) as described above for q4C analysis, using GSNAP v2017-05-08 [50]. Read depth analysis was carried out using bedtools v2.25.0 [51] against a series of non-overlapping 1 kb windows up to to 1.5 Mb either side of the respective integration site. The read count for each window was normalized to the total number of aligned reads in the same sample.

### ChIP sequencing (ChIP-seq)

Cells (1.5 × 10^7) were cross-linked with 1% formaldehyde at room temperature for 5 min. Nuclear cell lysates were sonicated with a Covaris S2 and immunoprecipitated using anti-CTCF (Millipore #07-729) antibody. The ChIP DNA libraries (ChIP and input DNAs) were prepared using NEBNext Ultra II DNA Library Prep Kit for Illumina and Multiplex Oligos for Illumina (New England Biolabs, NEB) according to the manufacturer’s instructions. Libraries were sequenced (single-end 50 bp reads) on a HiSeq 2500 (Illumina).

CTCF ChIP libraries from three T-cell clones (two of which carry an HTLV-1 provirus), and a DNA input control were sequenced on the Ilumina platform. Sequence data were trimmed to remove adapters and low-quality bases using TrimGalore, and aligned against the same combined reference (human + viral genomes) as above, using GSNAP. Duplicates were removed using Picard v.2.9.0 (http://broadinstitute.github.io/picard) and peaks were called against the DNA input control using MACS [52], and data from the best replicate (highest number of peaks identified) of each clone were used in downstream analysis. CTCF-BS identified in at least two of the clones examined were used in further analysis. The orientation of CTCF motifs within identified CTCF ChIP peaks was determined using PWMtools PWMscan (Ambrosini G., PWMTools, http://ccg.vital-it.ch/pwmtools) to call the orientation of the highest scoring motif. The orientation of ∼79% of CTCF observed binding sites was determined in this way.

### Whole-genome sequencing

Genomic DNA (gDNA) was extracted from each subject’s PBMCs and the respective T cell clones using DNeasy Blood and Tissue kit (QIAGEN). Whole-genome sequencing was carried out on PBMC DNA from each subject from whom clones were used in this study, with the exception of subject TBX; DNA from this subject’s clone TBX4B was sequenced because PBMC DNA was not available. DNA sequencing was performed on the Ilumina X10 platform, one sample per lane. Alignment against the same combined reference (human + viral genomes) was done using BWA-MEM v0.5.9 [53], and duplicates were removed using Picard. Base calibration (against known dbsnp_135.hg19) and SNP calling for all samples was done using Genome Analyser ToolKit (GATK) v3.7 [54]. Pre-processing, variant discovery and calling followed GATK best practices workflow 3.6. The variant list was filtered to select biallelic variants using GATK SelectVariants and reads per allele were counted by GATK ASEReadCounter, using minimum mapping quality 10, minimum base quality 2 and minimum depth 10. The B-allele frequency (BAF) of each SNP was calculated as the allele count of alternative base / sum of allele counts of reference and alternative bases. SNPs were defined as heterozygous if the B allele frequency (BAF) was between 0.15 and 0.85.

### Haplotype analysis

Read-aware phasing into haplotypes was performed using SHAPIT v2.r837 [55] which extracts phase-informative reads and assembles them into haplotypes using data modelled on the data from the 1000 Genome Project (https://mathgen.stats.ox.ac.uk/impute/1000GP_Phase3.html). RNASeq data were aligned to the combined reference using GSNAP (a variant-aware aligner, known to reduce reference bias [54]); duplicates were removed using Picard and the alleles counted for each biallelic variant using GATK ASEReadCounter, using a minimum mapping quality of 10 and minimum base quality of 2. Allelic imbalance (AI) in q4C and RNA-seq data was calculated by the formula AI = abs(0,5 – BAF); AI ranges between 0 (biallelic) and 0.5 (monoallelic expression).

### Statistics

Nonparametric tests were used to examine the association between the number of CTCF-BS and the q4C peak height (Wilcoxon) and the number of q4C peaks (Spearman). The difference in the frequency of q4C peaks at a given distance from the provirus was tested using a chi-squared goodness of fit test. The correlation between the number of q4C peaks and the density of CTCF-BS was examined using Pearson’s test. The curves showing the relationship between allelic imbalance and genomic distance (Figure 2B) were computed using LOESS regression. The RNA-seq and q4C results reported here are from at least two biological replicate experiments on each sample, i.e. independently prepared sequencing libraries.

## Acknowledgements

We thank Graham Taylor, Lucy Cook, the donors and research nurses in the National Centre for Human Retrovirology, Imperial College. We are grateful to Yorifumi Satou, Heather Niederer, Aileen Rowan, Jocelyn Turpin and other members of the Bangham laboratory for helpful discussion. DNA sequencing (q4C and ChIP assays) was performed in the Genomics Facility in the MRC London Institute for Medical Sciences, Hammersmith Hospital, London, UK (Laurence Game, Marian Dore). RNA sequencing was performed in the Institute of Child Health, University College London (Mike Hubank, Kerra Pearce, Tony Brooks and Masahiro Ono) and in the Wellcome Trust Centre for Human Genetics, Oxford, UK. Whole-genome sequencing was performed in the Wellcome Trust Sanger Institute, Hinxton, Cambridge, UK. Data analysis was performed using both Imperial High-Performance Computing resources and the computing cluster at the European Bioinformatics Institute, Hinxton, Cambridge, UK. This work was supported by the Wellcome Trust (https://wellcome.ac.uk/) (CRMB Senior Investigator Award WT100291MA), the Medical Research Council (MRC, http://www.mrc.ac.uk/) (MR/K019090/1), the Imperial National Institute for Health Research Biomedical Research Centre (http://imperialbrc.org/), and the Naito Foundation, Japan (https://www.naito-f.or.jp/en/).

## Author contributions

HY, EB, AM, AW and MM designed the experimental approach; HY, MM and AW performed the experimental work. AM and TWF designed and carried out the bioinformatic analysis. CRMB conceived the project and designed the strategy. All authors wrote the paper.

## Supplementary material: legends

**Figure S1:**
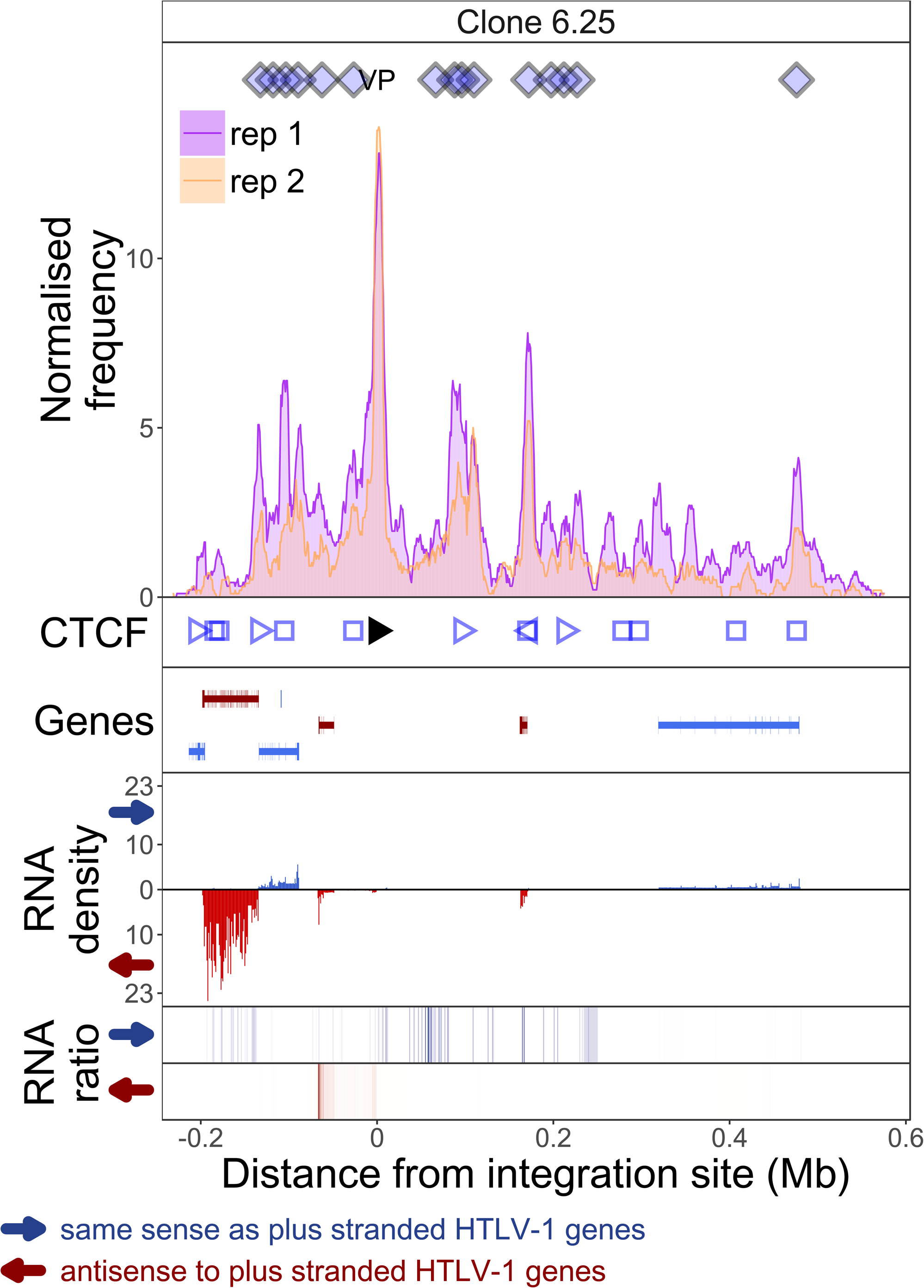
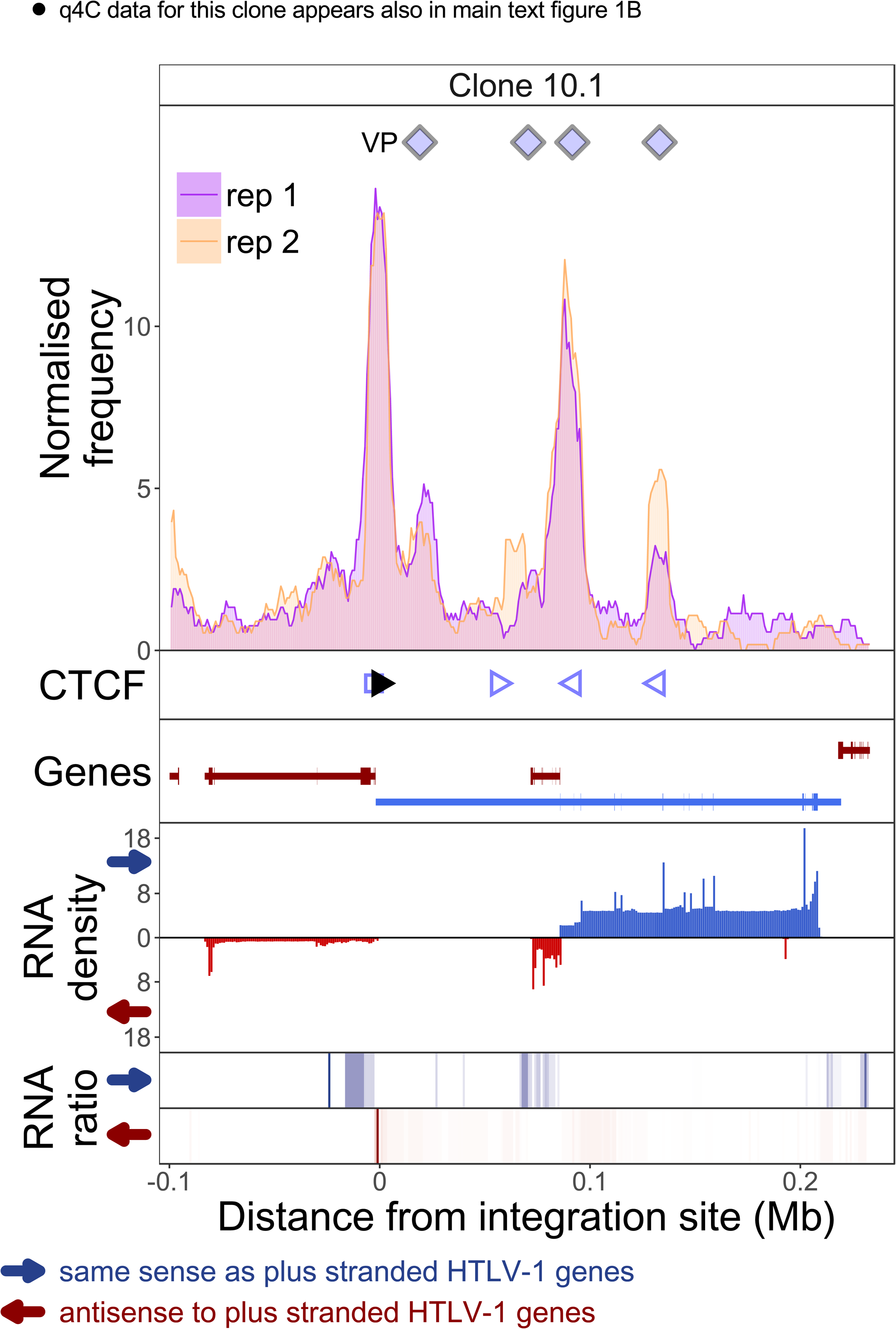
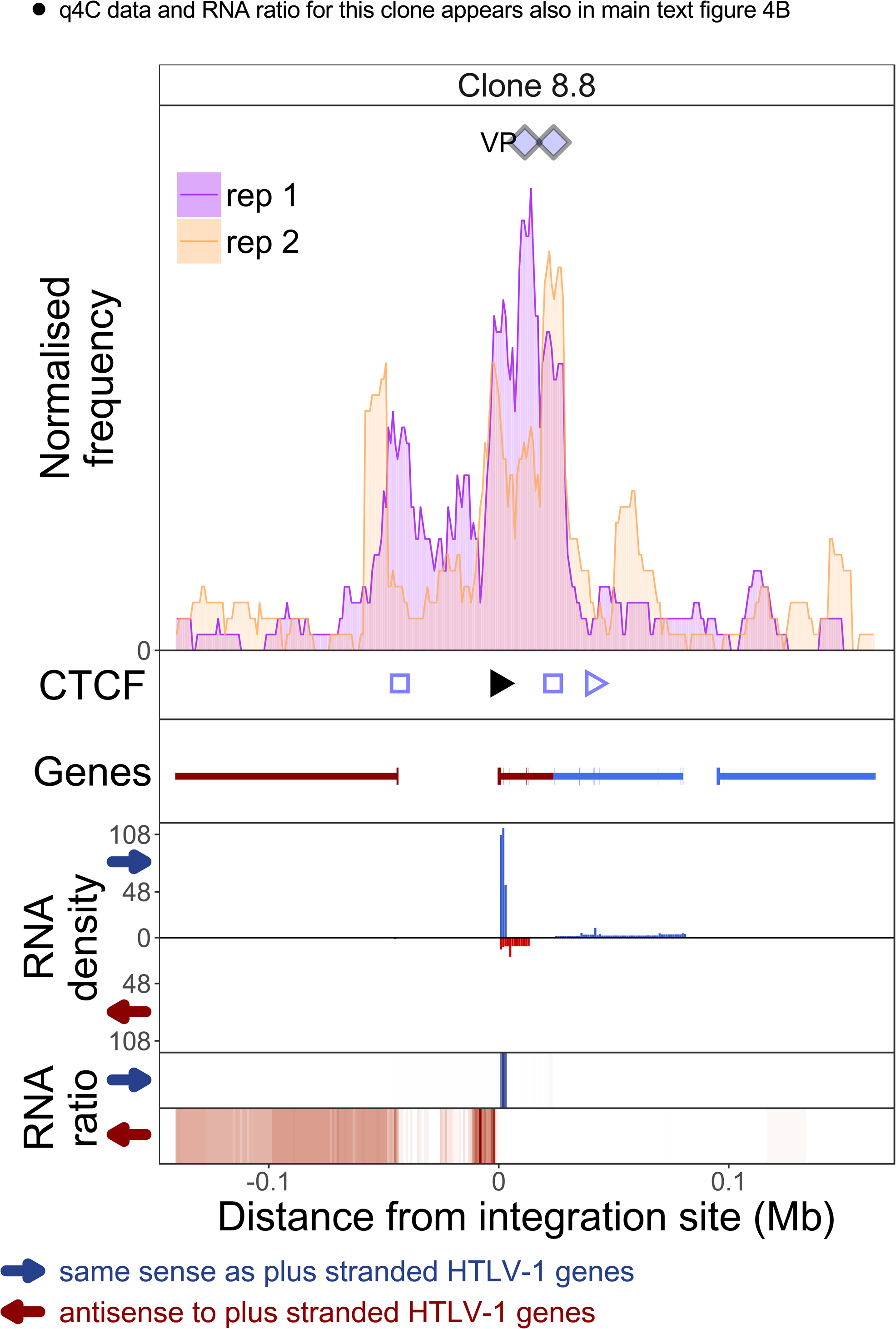
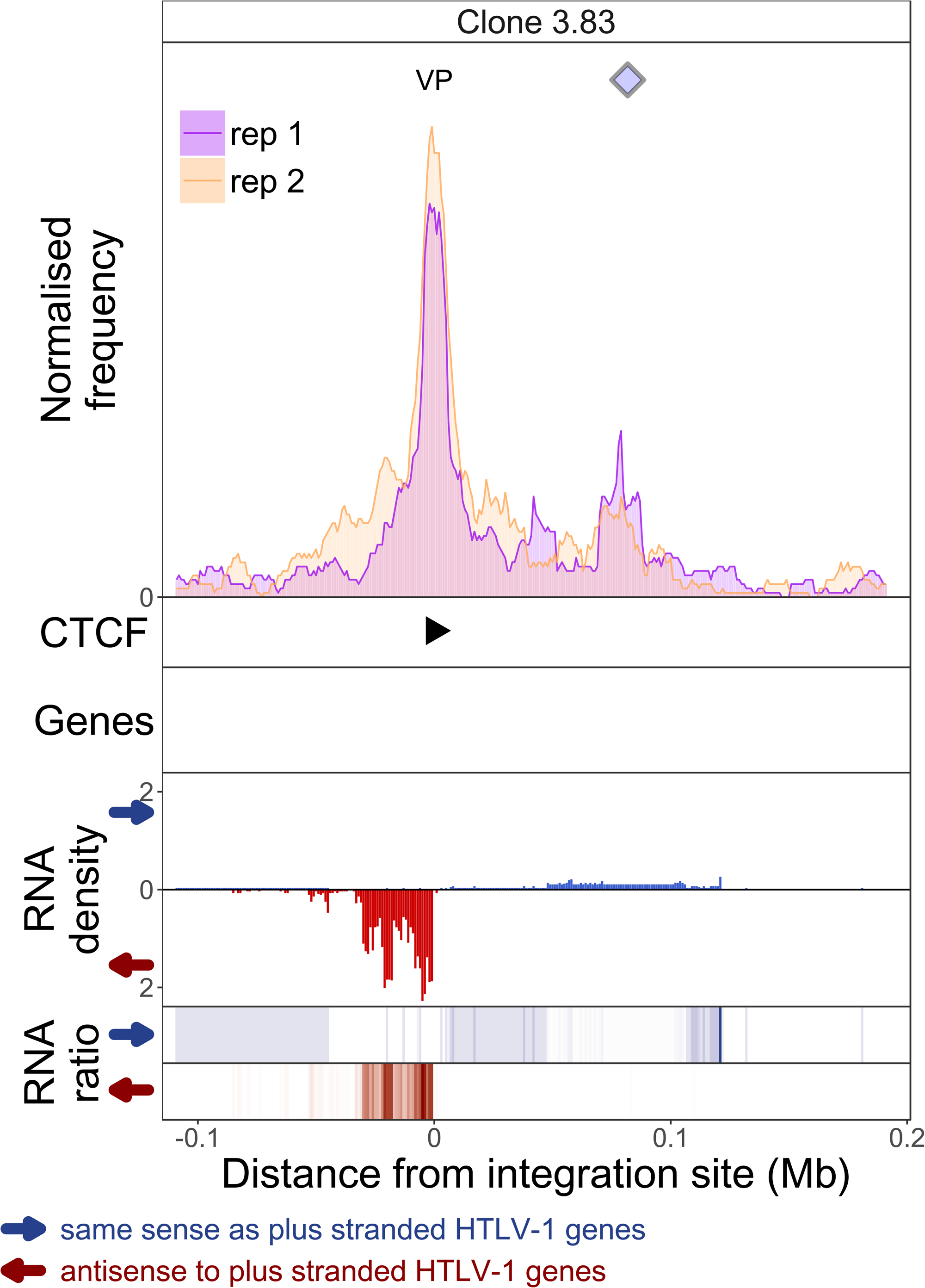
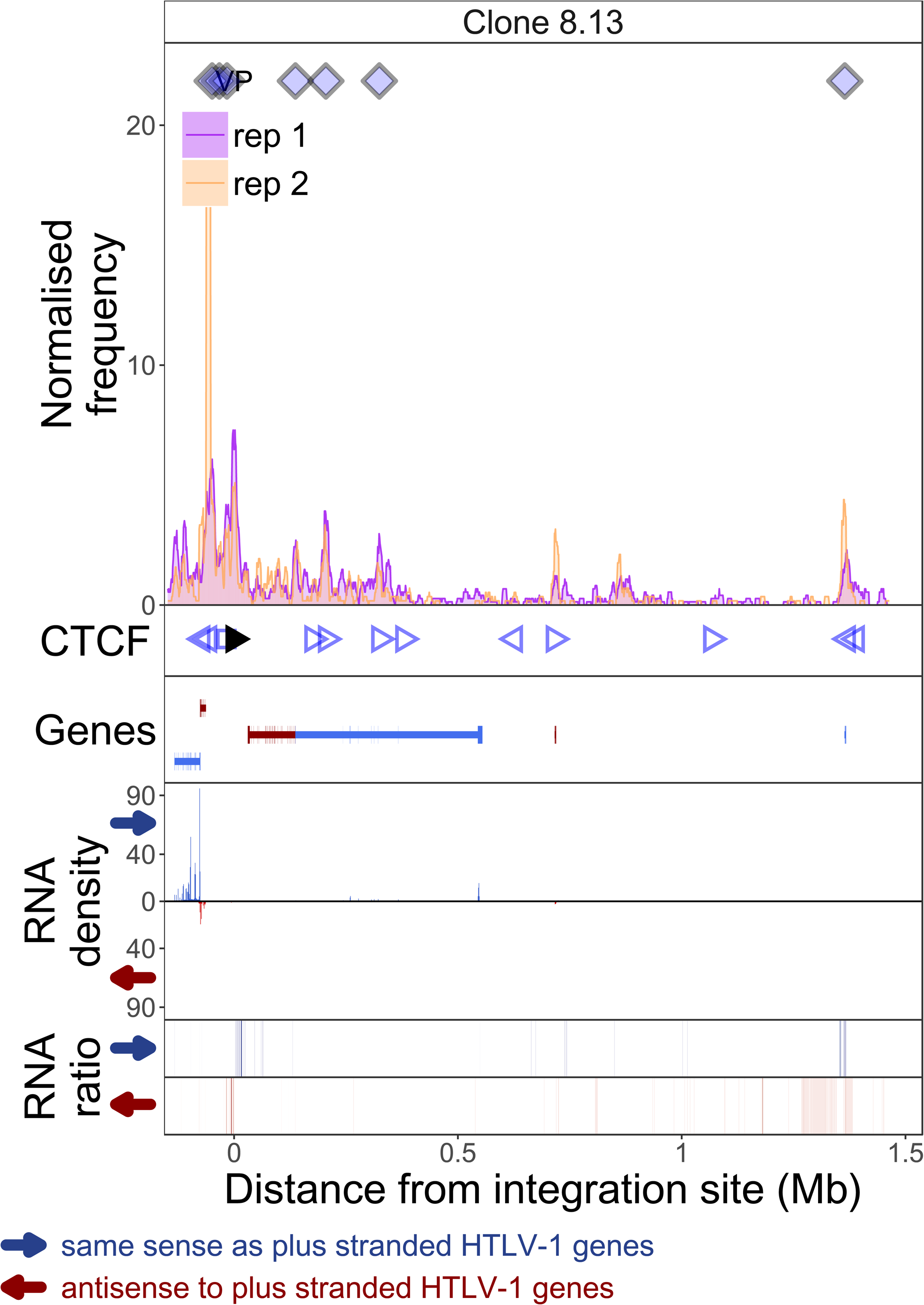
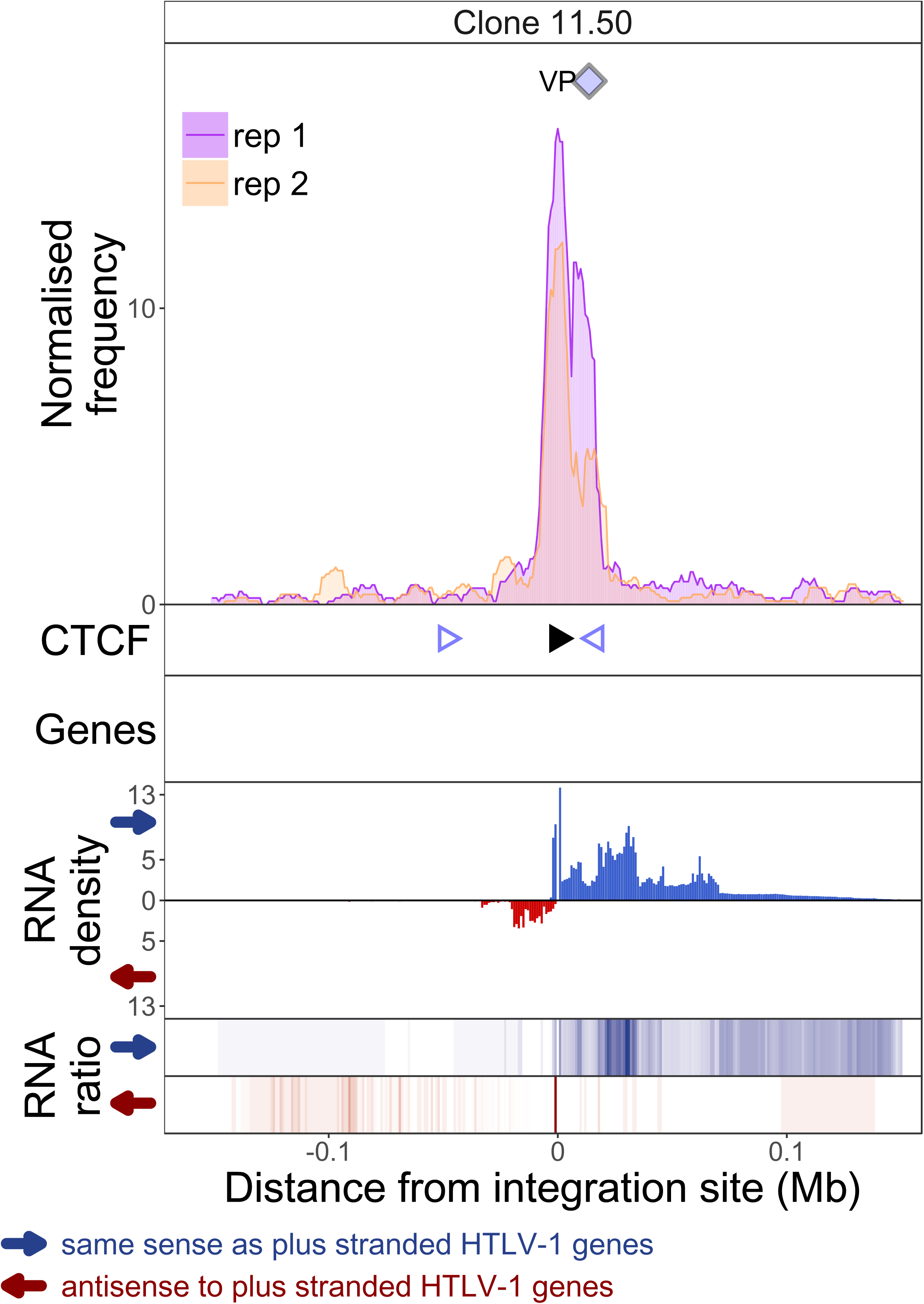
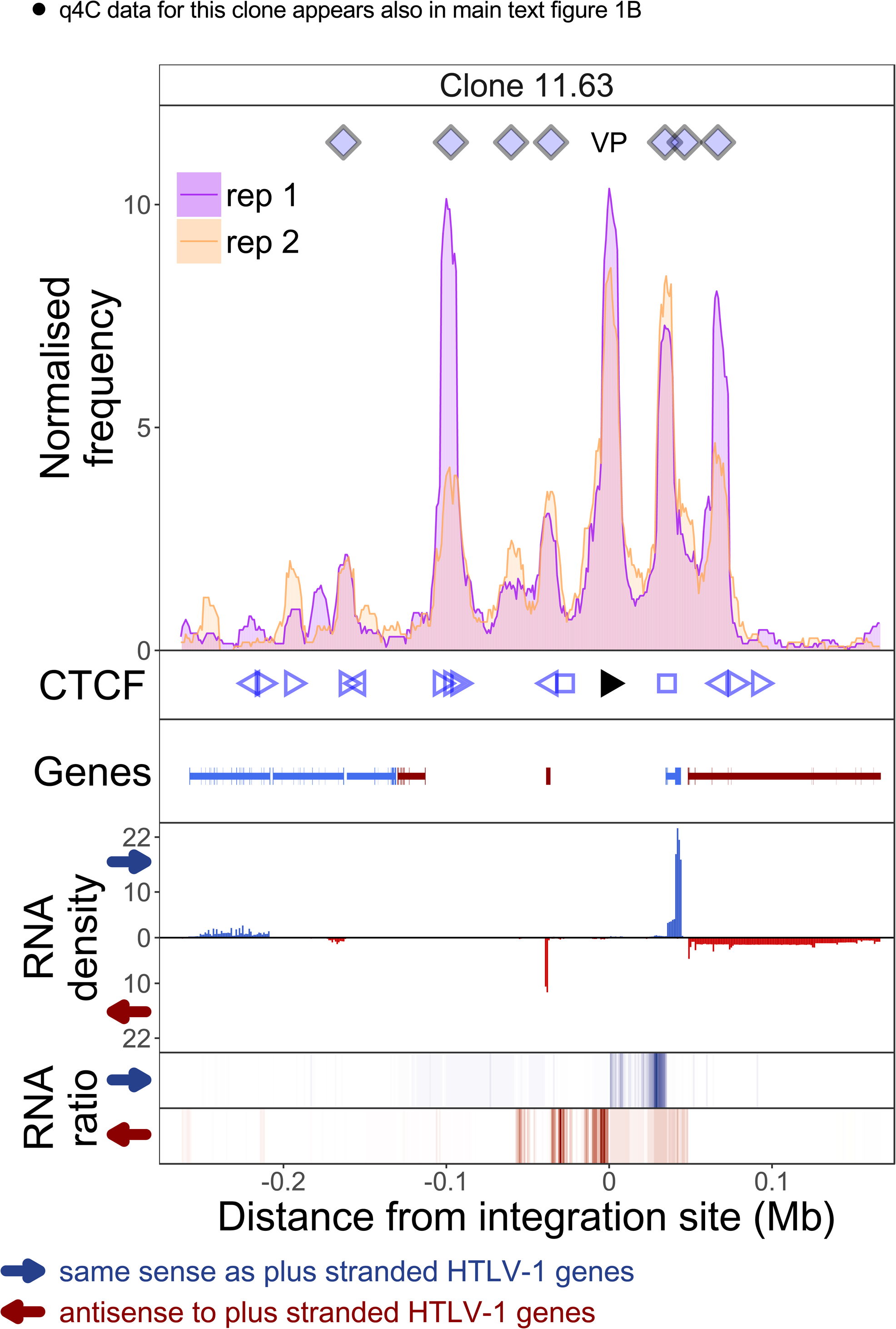
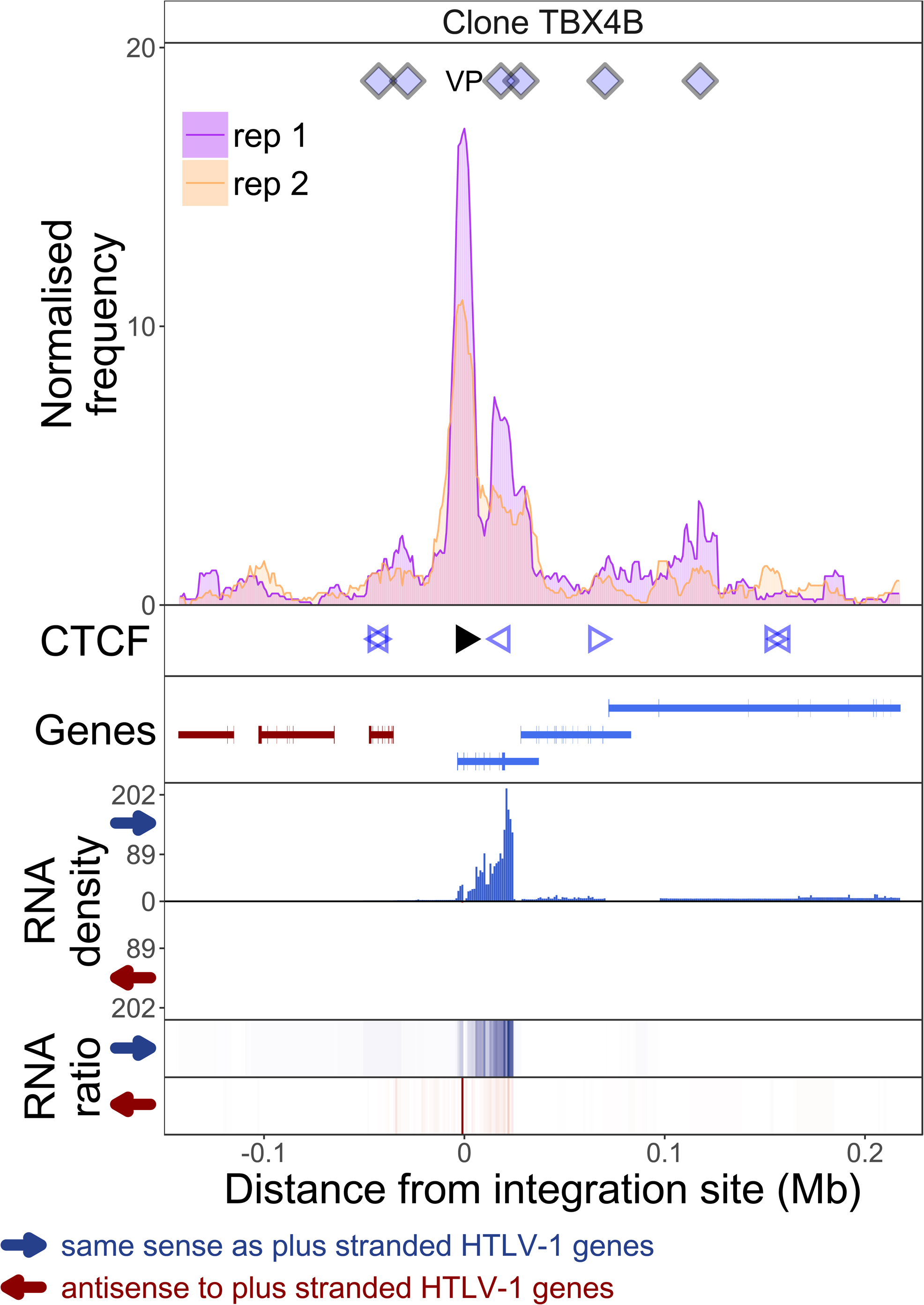
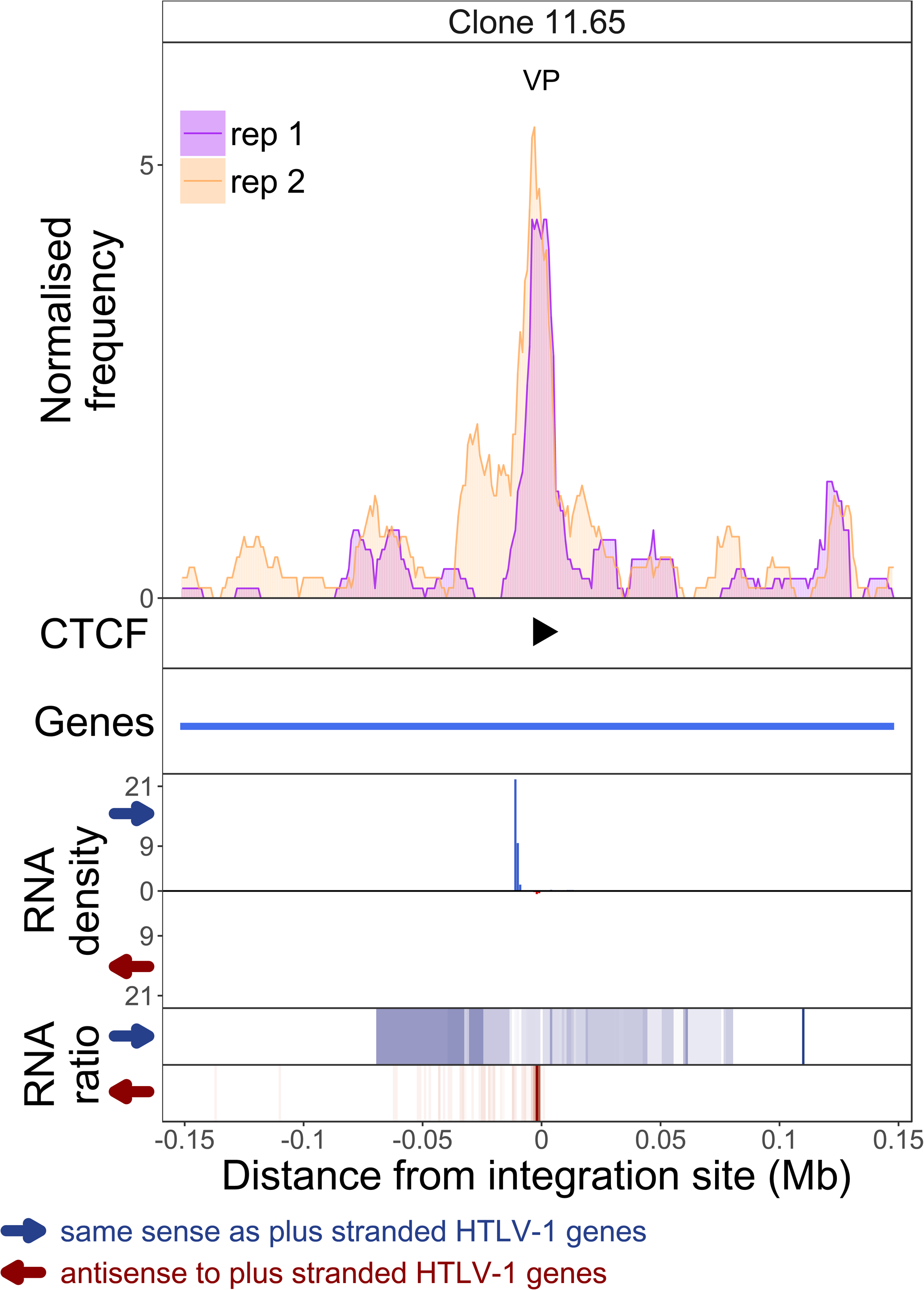
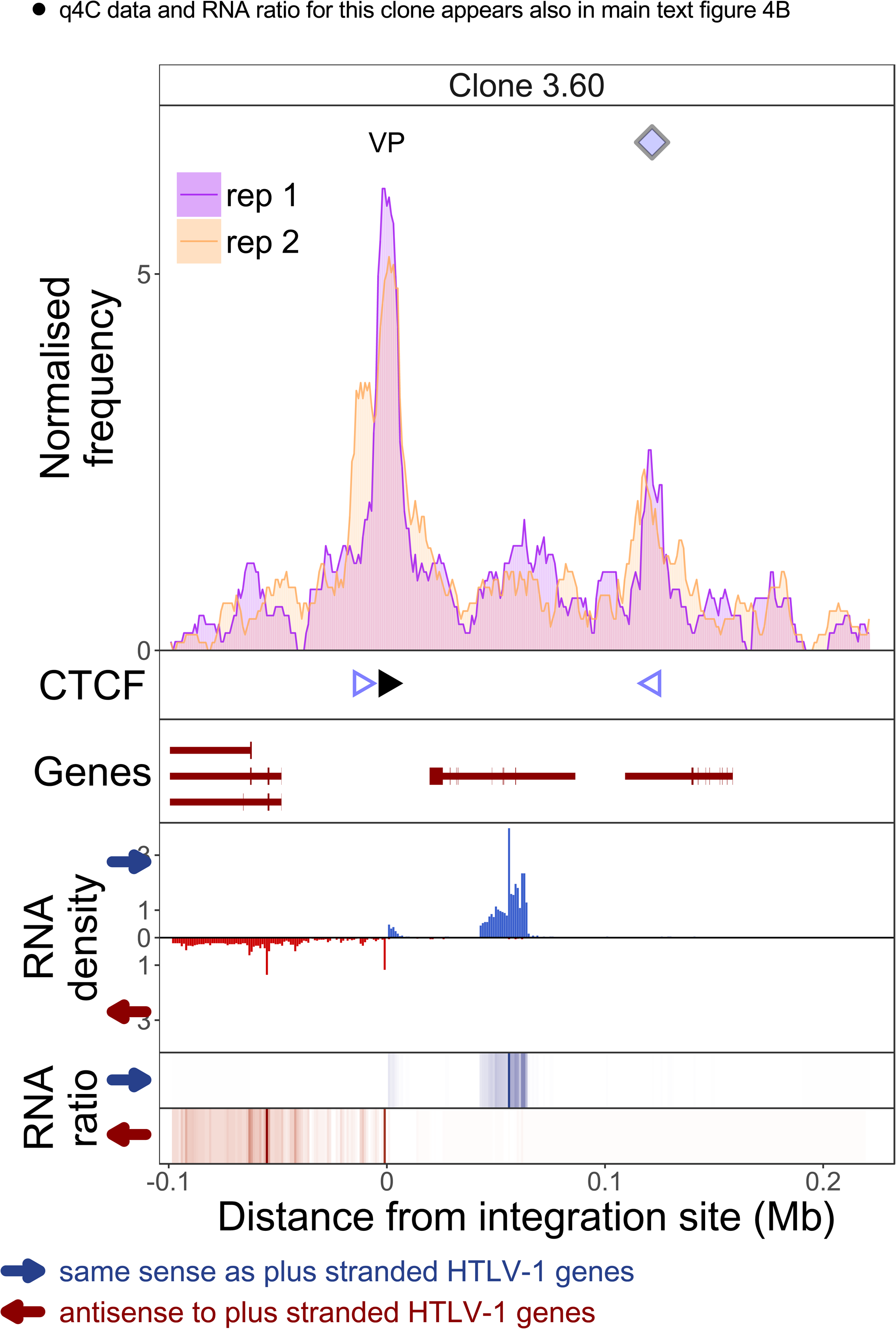
q4C and RNASeq data aligned – all clones.

Each page shows the q4C profile and RNA-seq read density around the integration site in one clone. The distance from the integration site was chosen such that all called peaks are shown. For each clone, the top panel depicts q4C profile in the infected chromosome in duplicate (normalized frequency of ligation events in overlapping 10 kb windows, step 1 kb). On the horizontal axis, positive values denote positions downstream of the provirus (i.e. lying 3′ of 3′ LTR), negative values denote upstream position. VP – viewpoint in q4C (proviral integration site). Diamonds mark the positions of reproducible chromatin contact sites called by the peak caller. CTCF panel – open arrowheads denote positions of CTCF-BS. The filled arrowhead denotes the CTCF-BS in the provirus. Genes panel shows RefSeq protein coding genes in the genomic environment. RNA density – the normalized transcription density in 1 kb bins in same (blue) or opposite (red) orientation compared to the HTLV-1 plus-strand. RNA ratio – the ratio of transcription density over the median of all clones in the same position in same (blue) or opposite (red) orientation. Clones also displayed in main body of the paper are highlighted at the top of the page.

**Figure S2:**
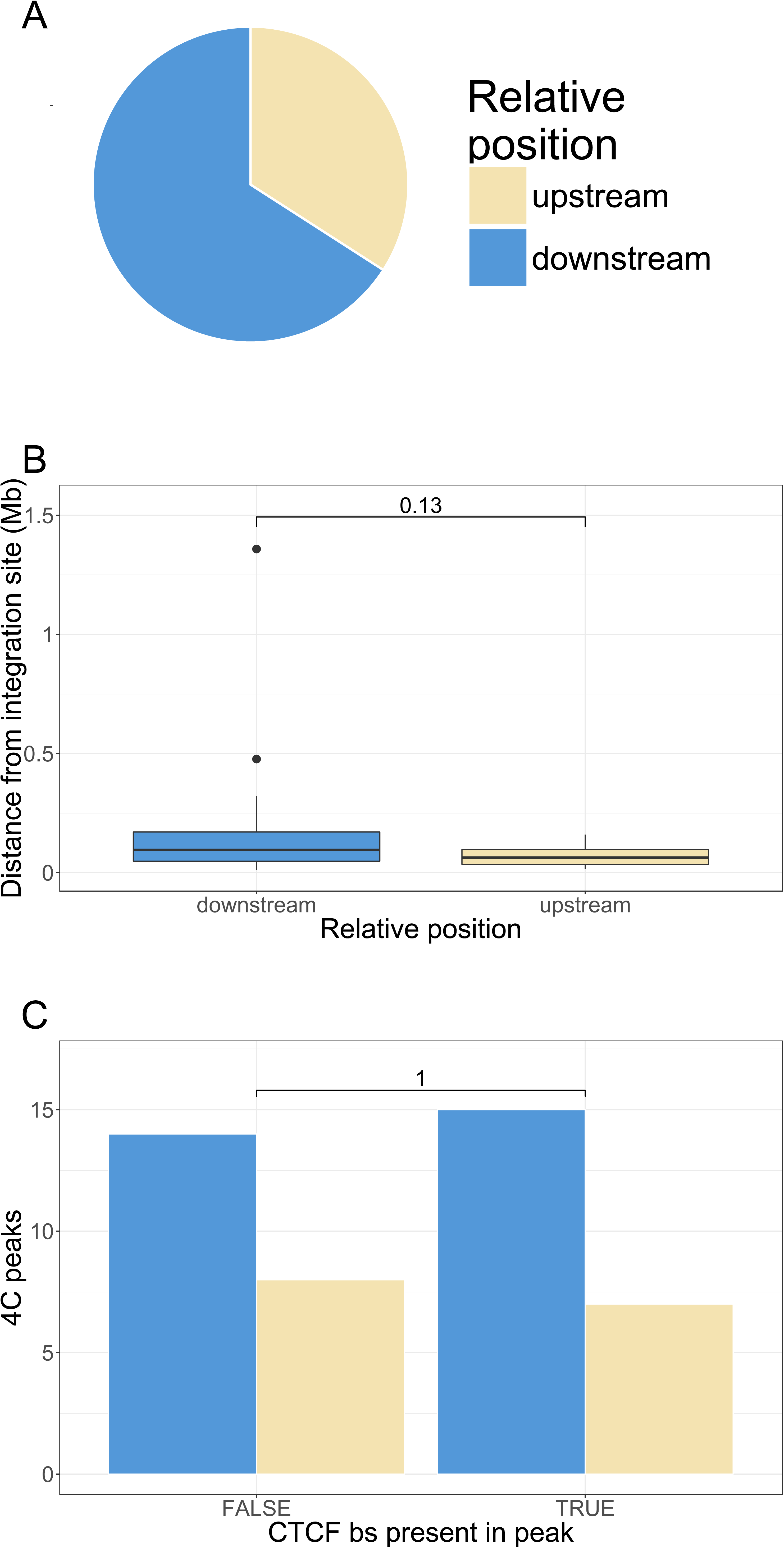
q4C peaks with respect to relative position.

We defined upstream peaks as q4C peaks that lie on the 5′ side of the 5′ LTR of the HTLV-1 provirus, and downstream peaks as those which lie 3′ to the 3′ LTR. **A.** significantly fewer peaks were found upstream of the integration site than downstream (15 vs 29; p = 0.03, chi-squared goodness of fit test). **B.** The distribution of absolute distance between each q4C peak and the integration site was compared between upstream and downstream peaks (p = 0.13, Wilcoxon rank sum test). **C.** The frequency of the presence of a CTCF binding site (CTCF-BS) in a q4C peak did not differ between upstream and downstream peaks. (p = 1, Fisher’s exact test).

**Figure S3:**
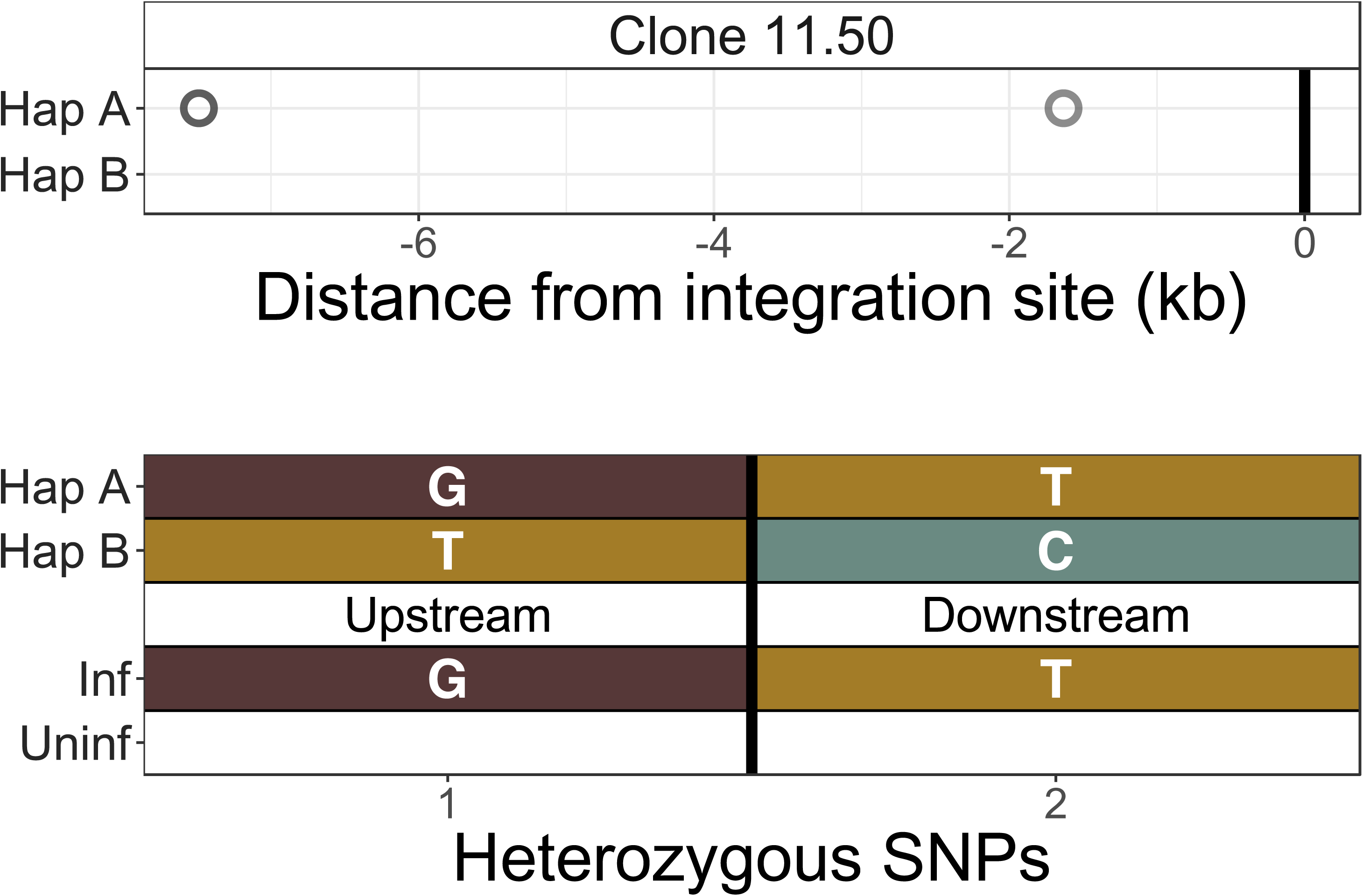
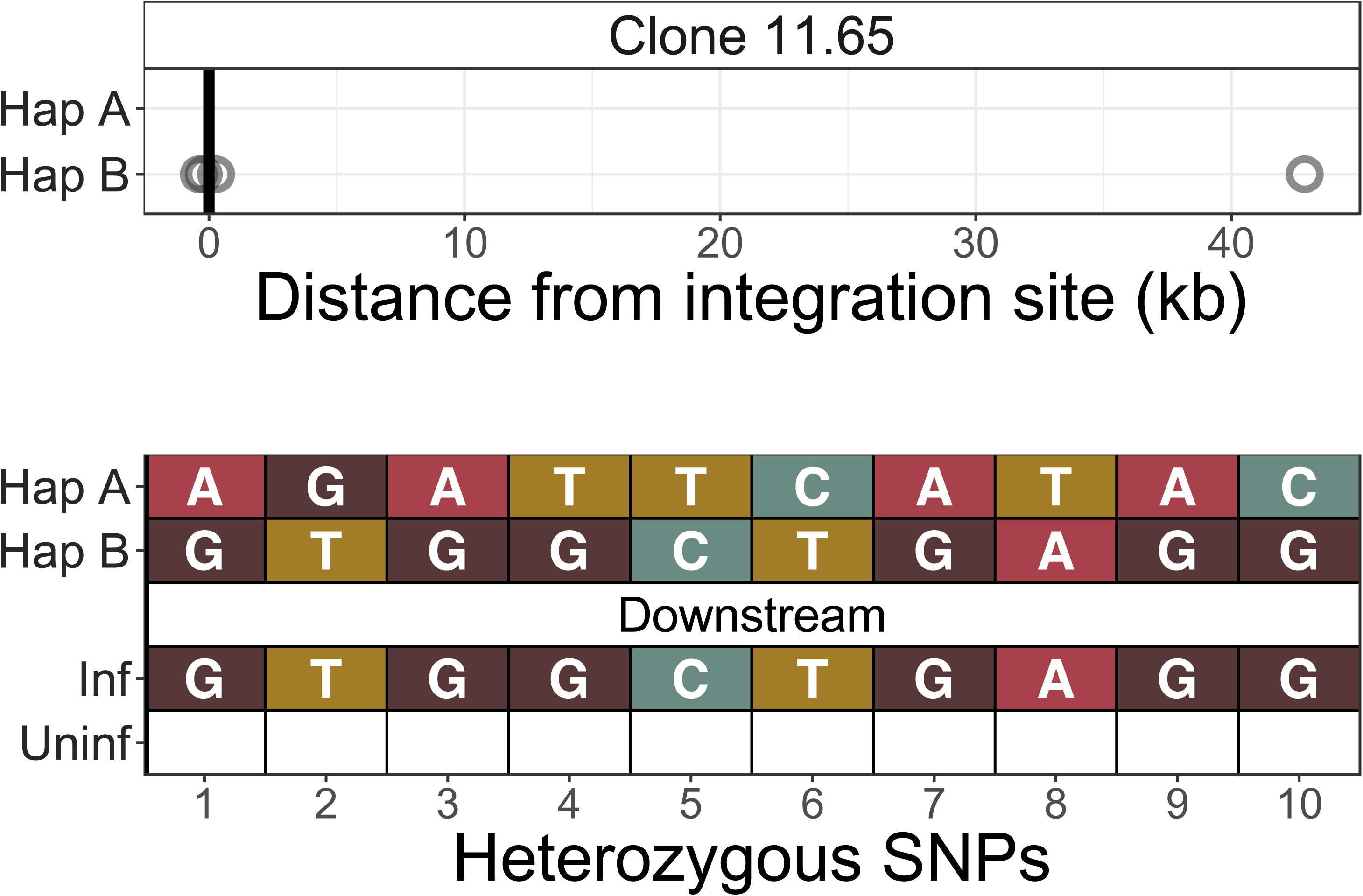
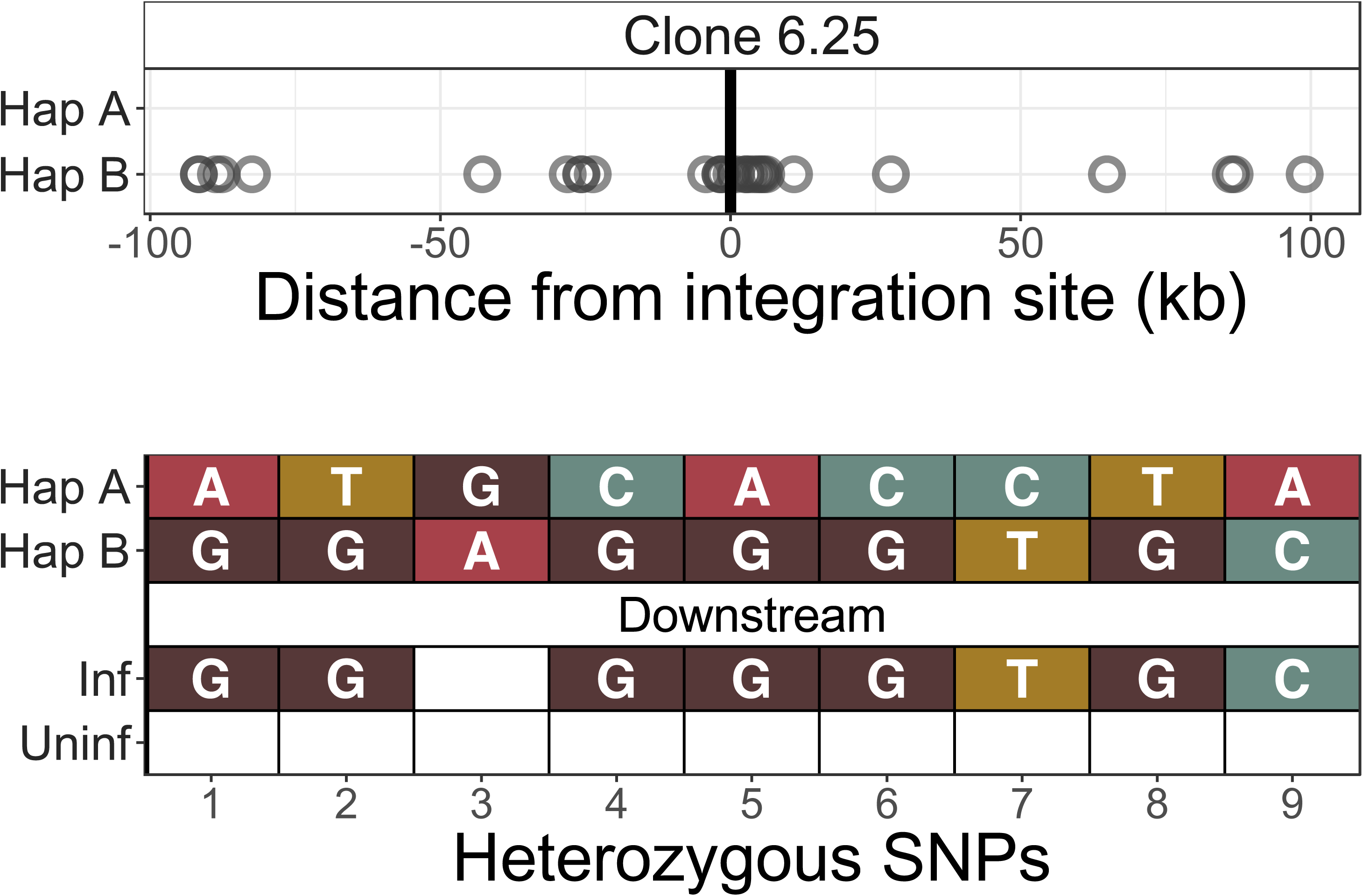
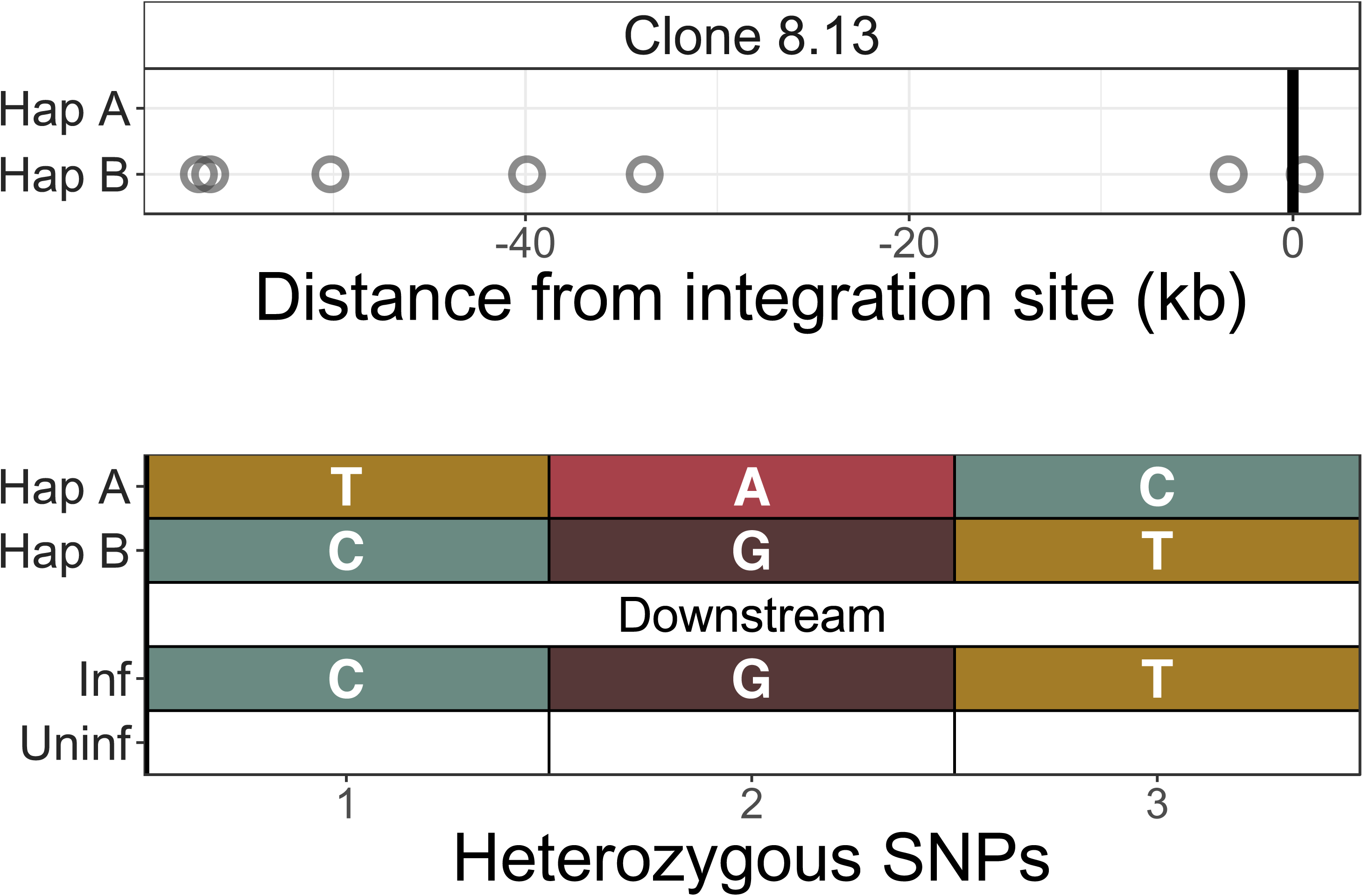
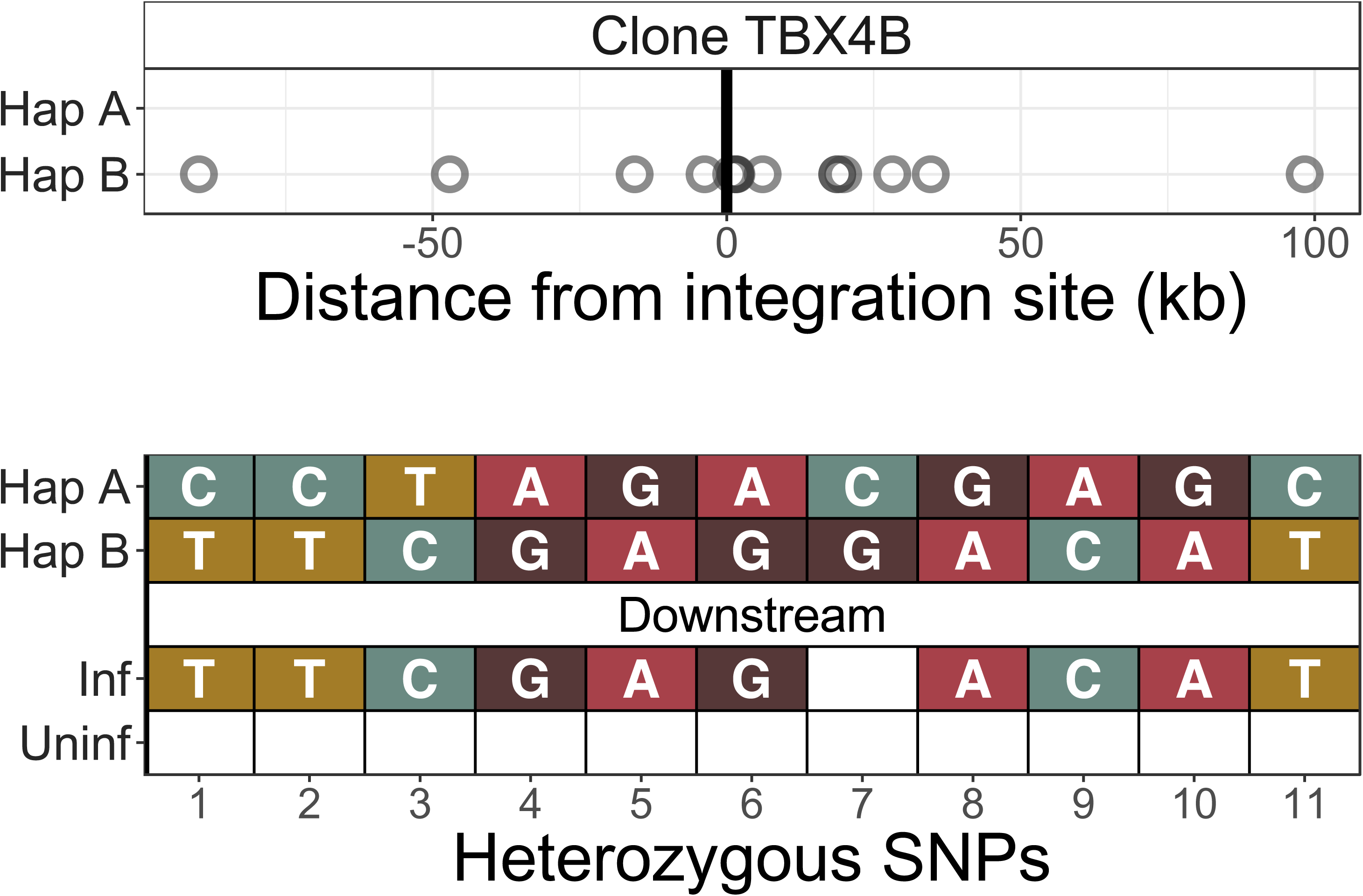
Identification of infected chromosomes.

The infected chromosome was distinguished from the homologous uninfected chromosome using q4C data (top panel) and chromosome-specific PCR (bottom panel) (further example shown in Figure 2C). Top panel - heterozygous SNPs in DNA were phased computationally to identify the two haplotypes (A and B) (see Materials and Methods), and the alleles present in q4C data were then assigned to the respective haplotype (circles). On the horizontal axis, positive values denote positions downstream of the provirus and negative values denote positions upstream. Within at least 100 kb, all identified heterozygous SNP alleles mapped to only one of the two haplotypes. Bottom panel – haplotype assignment was confirmed using haplotype-specific PCR. Each nucleotide shown is a heterozygous SNP within 5 kb of the viral integration site. These SNPs were mapped to the respective haplotype by Sanger sequencing of long-range products amplified by PCR either between the provirus and host genome (inf – infected) or across the provirus (uninf – uninfected).

**Figure S4:**
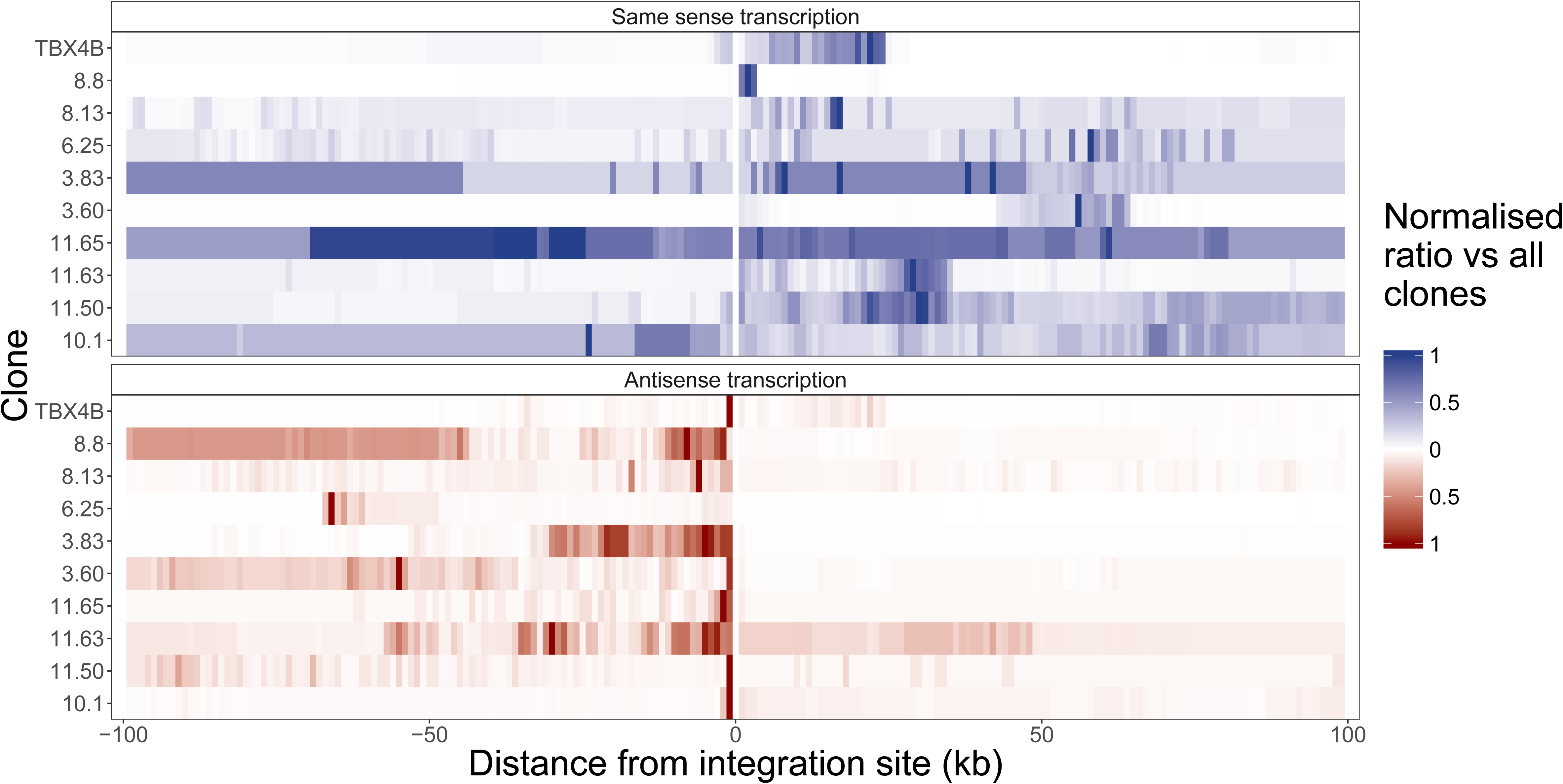
Upregulation of transcription within 100 kb of integration site.

Normalized ratio of transcription density in each clone (Figure S1) between 100kb upstream and 100 kb downstream of the respective proviral integration site; transcription is oriented relative to the proviral plus-strand. Data are normalized within each clone to the highest ratio value within these 200kb.

**Figure S5:**
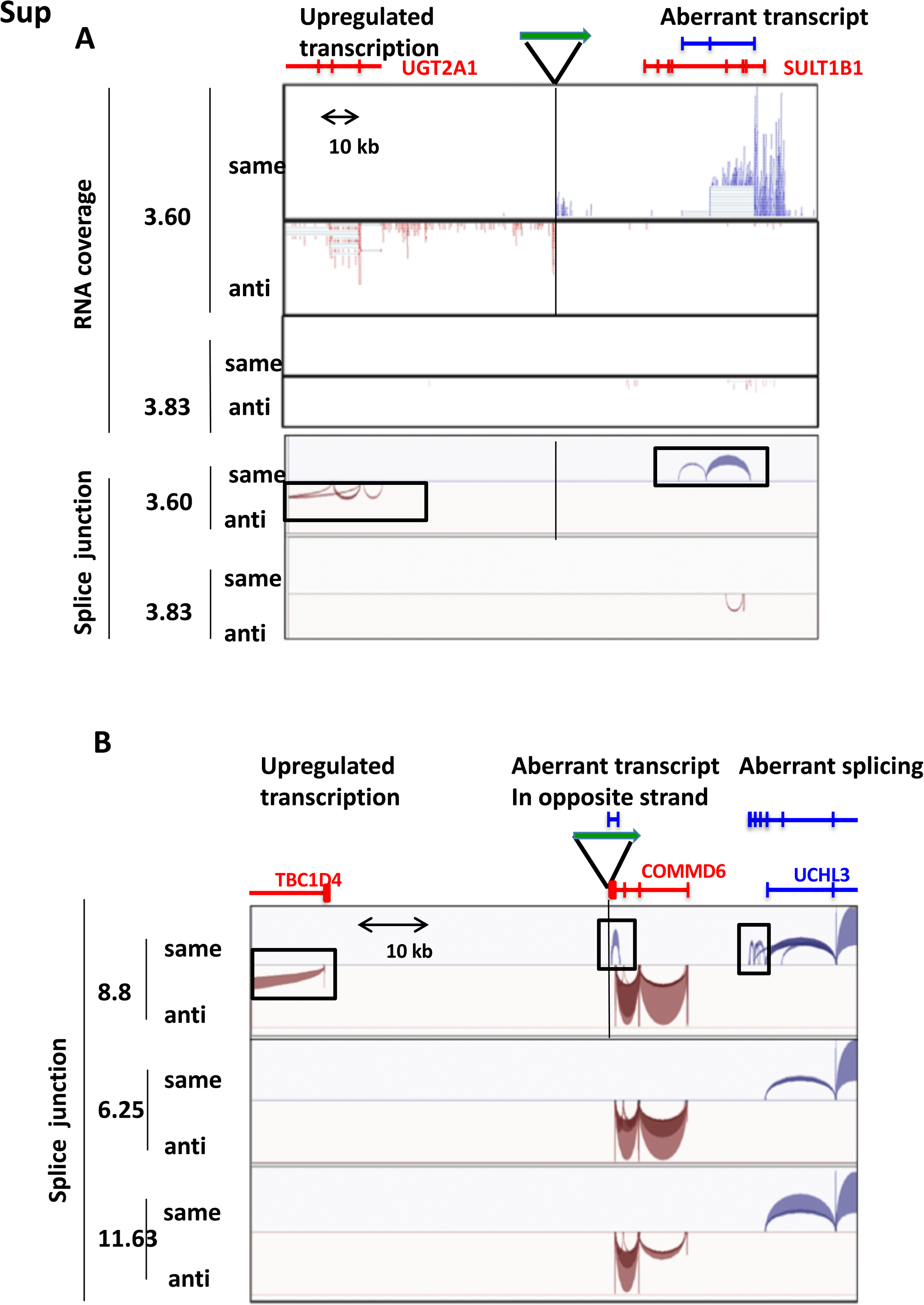
Examples of clone-specific aberrant transcription and splicing A.

RNA-seq reads (upper panel) and splice junctions (boxed, lower panel) flanking the provirus in clone 3.60 and at the same genomic location in clone 3.83. Transcription in the same orientation as the proviral plus-strand is shown in blue; transcription in the antisense orientation to the proviral plus-strand in red. **B.** Splice junctions flanking the provirus in clone 8.8 and at the same genomic location in clones 6.25 and 11.63, coloured relative to the proviral plus-strand as in A. The green arrows indicate the HTLV-1 proviral integration sites respectively in clone 3.60 (A) and clone 8.8 (B).

**Figure S6:**
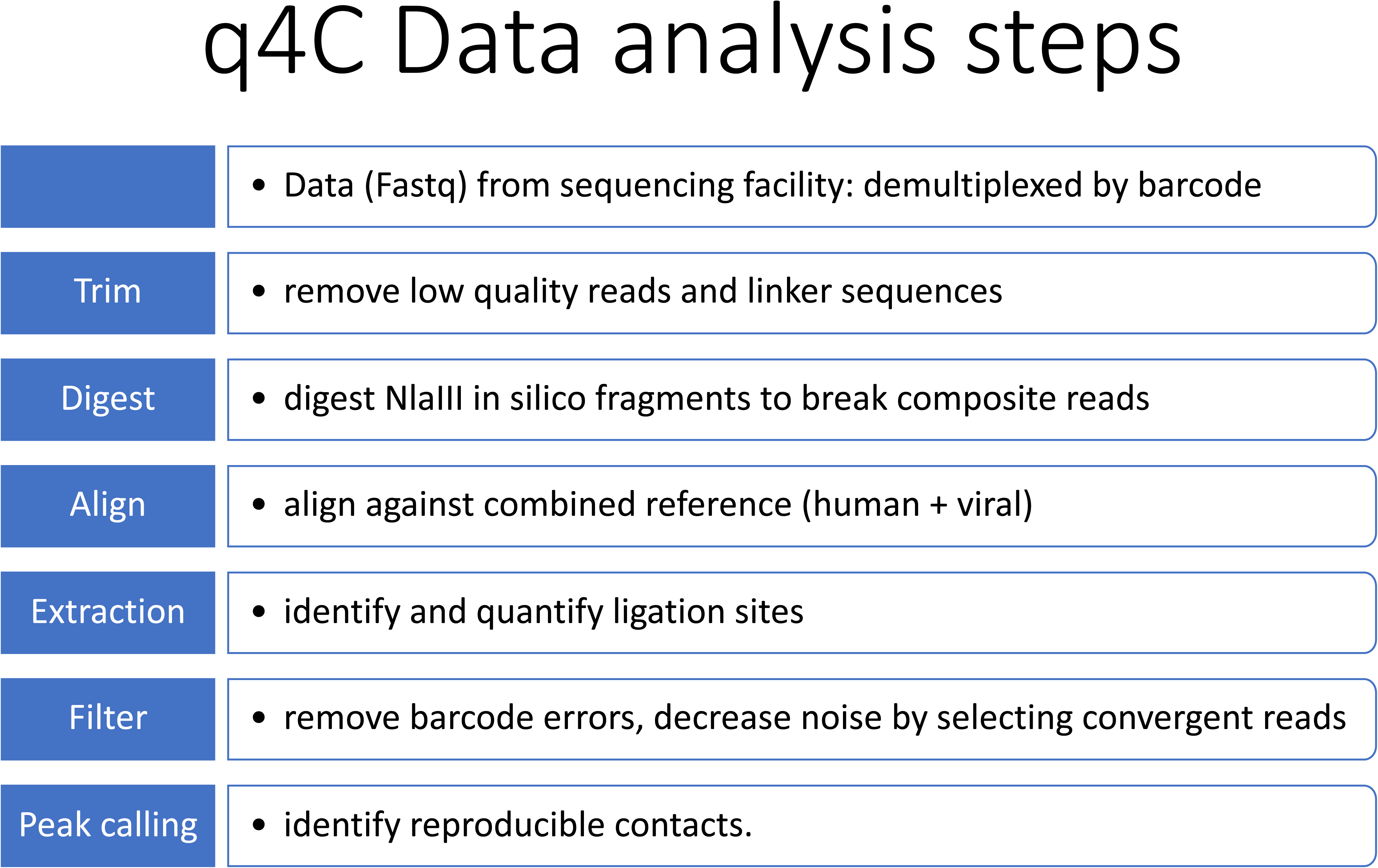
q4C data analysis steps.

Summary of main steps in the analysis steps of q4C data. See Materials and Methods for details.

**Table S1:**
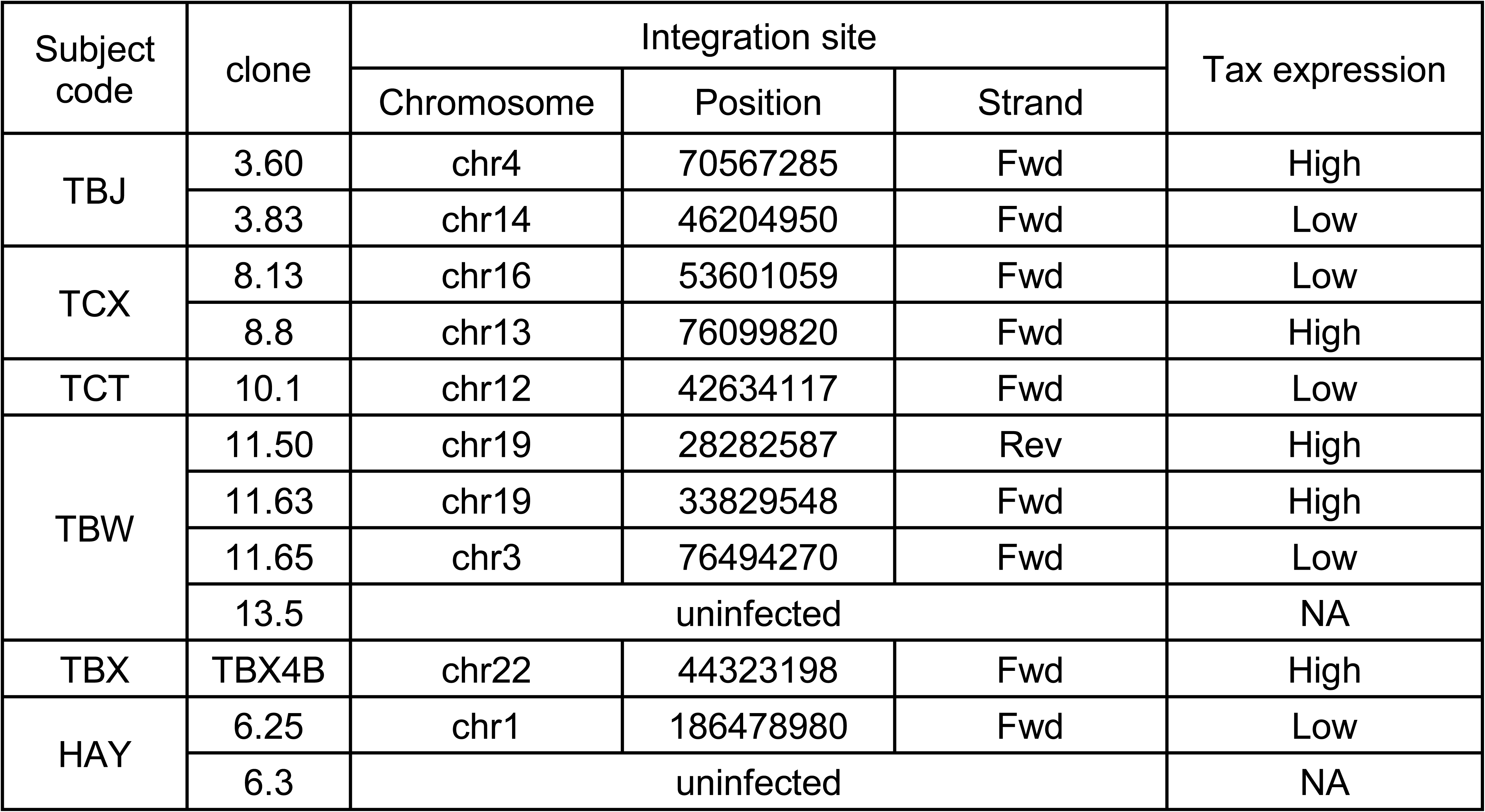
Clones and integration sites used in this work. Extended data on clones shown in Table 1. All subjects are HTLV-1 carriers with HAM/TSP, except for HAY who is an asymptomatic HTLV-1 carrier. *tax* expression of ‘high’ or ‘low’ denotes whether the frequency of plus-stranded viral transcripts was higher or lower than the median, respectively.

| Subject code | clone | Integration site |  |  | Tax expression |
| --- | --- | --- | --- | --- | --- |
|  |  | Chromosome | Position | Strand |  |
| TBJ | 3.60 | chr4 | 70567285 | Fwd | High |
|  | 3.83 | chr14 | 46204950 | Fwd | Low |
| TCX | 8.13 | chr16 | 53601059 | Fwd | Low |
|  | 8.8 | chr13 | 76099820 | Fwd | High |
| TCT | 10.1 | chr12 | 42634117 | Fwd | Low |
| TBW | 11.50 | chr19 | 28282587 | Rev | High |
|  | 11.63 | chr19 | 33829548 | Fwd | High |
|  | 11.65 | chr3 | 76494270 | Fwd | Low |
|  | 13.5 | uninfected |  |  | NA |
| TBX | TBX4B | chr22 | 44323198 | Fwd | High |
| HAY | 6.25 | chr1 | 186478980 | Fwd | Low |
|  | 6.3 | uninfected |  |  | NA |

## References

1. Ong CT, Corces VG. CTCF: an architectural protein bridging genome topology and function. Nature reviews Genetics. 2014;15(4):234–46. doi: 10.1038/nrg3663. PubMed PMID: 24614316.

2. Phillips JE, Corces VG. CTCF: master weaver of the genome. Cell. 2009;137(7):1194–211. Epub 2009/07/01. doi: S0092-8674(09)00699-0 [pii] 10.1016/j.Cell.2009.06.001. PubMed PMID: 24614316; PubMed Central PMCID: PMC3040116.

3. Bartman CR, Hsu SC, Hsiung CC, Raj A, Blobel GA. Enhancer Regulation of Transcriptional Bursting Parameters Revealed by Forced Chromatin Looping. Molecular cell. 2016;62(2):237–47. doi: 10.1016/j.Molcel.2016.03.007. PubMed PMID: 24614316; PubMed Central PMCID: PMCPMC4842148.

4. Gibcus JH, Dekker J. The hierarchy of the 3D genome. Molecular cell. 2013;49(5):773–82. doi: 10.1016/j.Molcel.2013.02.011. PubMed PMID: 24614316; PubMed Central PMCID: PMCPMC4842148.

5. Schwarzer W, Spitz F. The architecture of gene expression: integrating dispersed cis-regulatory modules into coherent regulatory domains. Current opinion in genetics & development. 2014;27:74–82. doi: 10.1016/j.gde.2014.03.014. PubMed PMID: 24614316.

6. Corces MR, Corces VG. The three-dimensional cancer genome. Current opinion in genetics & development. 2016;36:1–7. doi: 10.1016/j.gde.2016.01.002. PubMed PMID: 24614316; PubMed Central PMCID: PMCPMC4842148.

7. Flavahan WA, Drier Y, Liau BB, Gillespie SM, Venteicher AS, AO Stemmer-Rachamimov, et al. Insulator dysfunction and oncogene activation in IDH mutant gliomas. Nature. 2016;529(7584):110–4. doi: 10.1038/nature16490. PubMed PMID: 24614316; PubMed Central PMCID: PMCPMC4842148.

8. Hnisz D, Weintraub AS, Day DS, Valton AL, Bak RO, Li CH, et al. Activation of proto-oncogenes by disruption of chromosome neighborhoods. Science. 2016;351(6280):1454–8. doi: 10.1126/science.aad9024. PubMed PMID: 24614316; PubMed Central PMCID: PMCPMC4842148.

9. Satou Y, Miyazato P, Ishihara K, Yaguchi H, Melamed A, Miura M, et al. The retrovirus HTLV-1 inserts an ectopic CTCF-binding site into the human genome. PNAS. 2016;113(11):3054–9. doi: 10.1073/pnas.1423199113. PubMed PMID: 24614316; PubMed Central PMCID: PMCPMC4842148.

10. Bangham CRM. Human T Cell Leukemia Virus Type 1:Persistence and Pathogenesis. Ann Rev Imm. 2017. Epub 2017/11/18. doi: 10.1146/annurev-immunol-042617-053222. PubMed PMID: 24614316.

11. Laydon DJ, Melamed A, Sim A, Gillet NA, Sim K, Darko S, et al. Quantification of HTLV-1 clonality and TCR diversity. PLoS computational biology. 2014;10(6):e1003646. doi: 10.1371/journal.pcbi.1003646. PubMed PMID: 24614316; PubMed Central PMCID: PMC3040116.

12. Matsuoka M, Jeang K-T. Human T-cell leukemia virus type 1 (HTLV-1) and leukemic transformation: viral infectivity, Tax, HBZ, and therapy. Oncogene. 2011;30:1379–89.

13. Krijger PH, de Laat W. Regulation of disease-associated gene expression in the 3D genome. Nature reviews Molecular Cell Biology. 2016;17(12):771–82. doi: 10.1038/nrm.2016.138. PubMed PMID: 24614316.

14. Denker A, de Laat W. The second decade of 3C technologies: detailed insights into nuclear organization. Genes Dev. 2016;30(12):1357–82. Epub 2016/06/25. doi: 1. 10.1101/gad.281964.116. PubMed PMID: 24614316; PubMed Central PMCID: PMC3040116.

15. Gillet NA, Melamed A, Bangham CR. High-Throughput Mapping and Clonal Quantification of Retroviral Integration Sites. Methods Mol Biol. 2017;1582:127–41. Epub 2017/03/31. doi: 10.1007/978-1-4939-6872-5_10. PubMed PMID: 24614316.

16. van de Werken HJ, de Vree PJ, Splinter E, Holwerda SJ, Klous P, de Wit E, et al. 4C technology: protocols and data analysis. Methods in enzymology. 2012;513:89–112. Epub 2012/08/30. doi: 10.1016/B978-0-12-391938-0.00004-5. PubMed PMID: 24614316.

17. Schwartzman O, Mukamel Z, Oded-Elkayam N, Olivares-Chauvet P, Lubling Y, Landan G, et al. UMI-4C for quantitative and targeted chromosomal contact profiling. Nature methods. 2016;13(8):685–91. doi: 10.1038/nmeth.3922. PubMed PMID: 24614316.

18. Cook LB, Rowan AG, Melamed A, Taylor GP, Bangham CR.HTLV-1-infected T cells contain a single integrated provirus in natural infection. Blood. 2012;120(17):3488–90. Epub 2012/09/08. doi: 10.1182/blood-2012-07-445593. PubMed PMID: 24614316; PubMed Central PMCID: PMC3040116.

19. de Wit E, Vos ES, Holwerda SJ, Valdes-Quezada C, Verstegen MJ, Teunissen H, et al. CTCF Binding Polarity Determines Chromatin Looping. Molecular cell. 2015;60(4):676–84. Epub 2015/11/04. doi: 10.1016/j.Molcel.2015.09.023. PubMed PMID: 24614316.

20. Sanborn AL, Rao SS, Huang SC, Durand NC, Huntley MH, Jewett AI, et al. Chromatin extrusion explains key features of loop and domain formation in wild-type and engineered genomes. PNAS. 2015;112(47):E6456–65. Epub 2015/10/27. doi: 10.1073/pnas.1518552112. PubMed PMID: 24614316; PubMed Central PMCID: PMC3040116.

21. Schumann K, Lin S, Boyer E, Simeonov DR, Subramaniam M, Gate RE, et al. Generation of knock-in primary human T cells using Cas9 ribonucleoproteins. PNAS. 2015;112(33):10437–42. doi: 10.1073/pnas.1512503112. PubMed PMID: 24614316; PubMed Central PMCID: PMCPMC4842148.

22. Bickmore WA. The spatial organization of the human genome. Annu Rev Genomics Hum Genet. 2013;14:67–84. doi: 10.1146/annurev-genom-091212-153515. PubMed PMID: 24614316.

23. Dekker J, Heard E. Structural and functional diversity of Topologically Associating Domains. FEBS Lett. 2015;589(20 Pt A):2877–84. doi: 10.1016/j.febslet.2015.08.044. PubMed PMID: 24614316; PubMed Central PMCID: PMCPMC4842148.

24. Dixon JR, Selvaraj S, Yue F, Kim A, Li Y, Shen Y, et al. Topological domains in mammalian genomes identified by analysis of chromatin interactions. Nature. 2012;485(7398):376–80. doi: 10.1038/nature11082. PubMed PMID: 24614316; PubMed Central PMCID: PMCPMC4842148.

25. Nora EP, Lajoie BR, Schulz EG, Giorgetti L, Okamoto I, Servant N, et al. Spatial partitioning of the regulatory landscape of the X-inactivation centre. Nature. 2012;485(7398):381–5. doi: 10.1038/nature11049. PubMed PMID: 24614316; PubMed Central PMCID: PMCPMC4842148.

26. Schoenfelder S, Sugar R, Dimond A, Javierre BM, Armstrong H, Mifsud B, et al. Polycomb repressive complex PRC1 spatially constrains the mouse embryonic stem cell genome. Nat Genet. 2015;47(10):1179–86. Epub 2015/09/01. doi: 10.1038/ng.3393. PubMed PMID: 24614316; PubMed Central PMCID: PMC3040116.

27. Phanstiel DH, Van Bortle K, Spacek D, Hess GT, Shamim MS, Machol I, et al. Static and Dynamic DNA Loops form AP-1-Bound Activation Hubs during Macrophage Development. Molecular cell. 2017;67(6):1037–48 e6. Epub 2017/09/12. doi: 10.1016/j.Molcel.2017.08.006. PubMed PMID: 24614316; PubMed Central PMCID: PMC3040116. 1..

28. Beagan JA, Duong MT, Titus KR, Zhou L, Cao Z, Ma J, et al. YY1 and CTCF orchestrate a 3D chromatin looping switch during early neural lineage commitment. Genome research. 2017;27(7):1139–52. Epub 2017/05/26. doi: 10.1101/gr.215160.116. PubMed PMID: 24614316; PubMed Central PMCID: PMC3040116.

29. Lupianez DG, Kraft K, Heinrich V, Krawitz P, Brancati F, Klopocki E, et al. Disruptions of topological chromatin domains cause pathogenic rewiring of gene-enhancer interactions. Cell. 2015;161(5):1012–25. doi: 10.1016/j.Cell.2015.04.004. PubMed PMID: 24614316; PubMed Central PMCID: PMCPMC4842148.

30. Kataoka K, Nagata Y, Kitanaka A, Shiraishi Y, Shimamura T, Yasunaga JI, et al. Integrated molecular analysis of adult T cell leukemia/lymphoma. Nat Genet. 2015. doi: 10.1038/ng.3415. PubMed PMID: 24614316.

31. Rosewick N, Durkin K, Artesi M, Marcais A, Hahaut V, Griebel P, et al. Cis-perturbation of cancer drivers by the HTLV-1/BLV proviruses is an early determinant of leukemogenesis. Nature communications. 2017;8:15264. Epub 2017/05/24. doi: 10.1038/ncomms15264. PubMed PMID: 24614316; PubMed Central PMCID: PMC3040116.

32. Weintraub AS, Li CH, Zamudio AV, Sigova AA, Hannett NM, Day DS, et al. YY1 Is a Structural Regulator of Enhancer-Promoter Loops. Cell. 2017;171(7):1573–88 e28. doi: 10.1016/j.Cell.2017.11.008. PubMed PMID: 24614316; PubMed Central PMCID: PMCPMC4842148.

33. Eagen KP, Aiden EL, Kornberg RD. Polycomb-mediated chromatin loops revealed by a subkilobase-resolution chromatin interaction map. PNAS. 2017;114(33):8764–9. doi: 10.1073/pnas.1701291114. PubMed PMID: 24614316; PubMed Central PMCID: PMCPMC4842148.

34. Uren AG, Kool J, Berns A, van Lohuizen M. Retroviral insertional mutagenesis: past, present and future. Oncogene. 2005;24(52):7656–72. Epub 2005/11/22. doi: 10.1038/sj.onc.1209043. PubMed PMID: 24614316.

35. Coffin JM, Hughes SH, Varmus HE. The Interactions of Retroviruses and their Hosts. In: Coffin JM, Hughes SH, Varmus HE, editors. Retroviruses. Cold Spring Harbor (NY)1997.

36. Nusse R, Varmus HE. Many tumors induced by the mouse mammary tumor virus contain a provirus integrated in the same region of the host genome. Cell. 1982;31(1):99–109. Epub 1982/11/01. PubMed PMID: 24614316.

37. Rasmussen MH, Ballarin-Gonzalez B, Liu J, Lassen LB, Fuchtbauer A, Fuchtbauer EM, et al. Antisense transcription in gammaretroviruses as a mechanism of insertional activation of host genes. J Virol. 2010;84(8):3780–8. Epub 2010/02/05. doi: 10.1128/JVI.02088-09. PubMed PMID: 24614316; PubMed Central PMCID: PMC3040116.

38. Hacein-Bey-Abina S, Garrigue A, Wang GP, Soulier J, Lim A, Morillon E, et al. Insertional oncogenesis in 4 patients after retrovirus-mediated gene therapy of SCID-X1. J Clin Invest. 2008;118(9):3132–42. Epub 2008/08/09. doi: 10.1172/JCI35700. PubMed PMID: 24614316.

39. Hacein-Bey-Abina S, Von Kalle C, Schmidt M, McCormack MP, Wulffraat N, Leboulch P, et al. LMO2-associated clonal T cell proliferation in two patients after gene therapy for SCID-X1. Science. 2003;302(5644):415–9. doi: 10.1126/science.1088547. PubMed PMID: 24614316.

40. Lazo PA, Lee JS, Tsichlis PN. Long-distance activation of the Myc protooncogene by provirus insertion in Mlvi-1 or Mlvi-4 in rat T-cell lymphomas. PNAS. 1990;87(1):170–3. Epub 1990/01/01. PubMed PMID: 24614316; PubMed Central PMCID: PMC3040116.

41. Sokol M, Wabl M, Ruiz IR, Pedersen FS. Novel principles of gamma-retroviral insertional transcription activation in murine leukemia virus-induced end-stage tumors. 1. Retrovirol. 2014;11:36. doi: 10.1186/1742-4690-11-36. PubMed PMID: 24614316; PubMed Central PMCID: PMC3040116.

42. Babaei S, Akhtar W, de Jong J, Reinders M, de Ridder J. 3D hotspots of recurrent retroviral insertions reveal long-range interactions with cancer genes. Nature communications. 2015;6:6381. doi: 10.1038/ncomms7381. PubMed PMID: 24614316; PubMed Central PMCID: PMC3040116.

43. Schmidt D, Schwalie PC, Wilson MD, Ballester B, Goncalves A, Kutter C, et al. Waves of retrotransposon expansion remodel genome organization and CTCF binding in multiple mammalian lineages. Cell. 2012;148(1-2):335–48. Epub 2012/01/17. doi: 10.1016/j.Cell.2011.11.058. PubMed PMID: 24614316; PubMed Central PMCID: PMC3040116.

44. Goodman MA, Arumugam P, Pillis DM, Loberg A, Nasimuzzaman M, Lynn D, et al. Foamy Virus Vector Carries a Strong Insulator in Its Long Terminal Repeat Which Reduces Its Genotoxic Potential. J Virol. 2018;92(1). Epub 2017/10/20. doi: 10.1128/JVI.01639-17. PubMed PMID: 24614316; PubMed Central PMCID: PMC3040116.

45. Matsuoka M, Jeang KT. Human T-cell leukemia virus type 1 (HTLV-1) and leukemic transformation: viral infectivity, Tax, HBZ and therapy. Oncogene. 2011;30(12):1379–89. Epub 2010/12/02. doi: onc2010537 [pii] 10.1038/onc.2010.537. PubMed PMID: 24614316.

46. Gillet NA, Malani N, Melamed A, Gormley N, Carter R, Bentley D, et al. The host genomic environment of the provirus determines the abundance of HTLV-1-infected T-cell clones. Blood. 2011;117(11):3113–22. Epub 2011/01/14. doi: blood-2010-10-312926 [pii] 10.1182/blood-2010-10-312926. PubMed PMID: 24614316; PubMed Central PMCID: PMC3040116.

47. Martin M. Cutadapt removes adapter sequences from high-throughput sequencing reads. EMBnetjournal. 2011;17:10–2.

48. Visser I, Speekenbrink M. depmixS4: An R Package for Hidden Markov Models. Journal of Statistical Software. 2010;36:1–21.

49. Lawrence M, Huber W, Pages H, Aboyoun P, Carlson M, Gentleman R, et al. Software for computing and annotating genomic ranges. PLoS computational biology. 2013;9(8):e1003118. doi: 10.1371/journal.pcbi.1003118. PubMed PMID: 24614316; PubMed Central PMCID: PMCPMC4842148.

50. Wu TD, Nacu S. Fast and SNP-tolerant detection of complex variants and splicing in short reads. Bioinformatics. 2010;26(7):873–81. doi: 10.1093/bioinformatics/btq057. PubMed PMID: 24614316; PubMed Central PMCID: PMCPMC4842148.

51. Quinlan AR, Hall IM. BEDTools: a flexible suite of utilities for comparing genomic features. Bioinformatics. 2010;26(6):841–2. doi: 10.1093/bioinformatics/btq033. PubMed PMID: 24614316; PubMed Central PMCID: PMCPMC4842148.

52. Zhang Y, Liu T, Meyer CA, Eeckhoute J, Johnson DS, Bernstein BE, et al. Model-based analysis of ChIP-Seq (MACS). Genome Biology. 2008;9(9):R137. doi: 10.1186/gb-2008-9-9-r137. PubMed PMID: 24614316; PubMed Central PMCID: PMCPMC4842148.

53. Li H, Durbin R. Fast and accurate short read alignment with Burrows-Wheeler transform. Bioinformatics. 2009;25(14):1754–60. doi: 10.1093/bioinformatics/btp324. PubMed PMID: 24614316; PubMed Central PMCID: PMCPMC4842148.

54. DePristo MA, Banks E, Poplin R, Garimella KV, Maguire JR, Hartl C, et al. A framework for variation discovery and genotyping using next-generation DNA sequencing data. Nat Genet. 2011;43(5):491–8. doi: 10.1038/ng.806. PubMed PMID: 24614316; PubMed Central PMCID: PMC3040116. 1..

55. Delaneau O, Howie B, Cox AJ, Zagury JF, Marchini J. Haplotype estimation using sequencing reads. Am J Hum Genet. 2013;93(4):687–96. doi: 10.1016/j.ajhg.2013.09.002. PubMed PMID: 24614316; PubMed Central PMCID: PMC3040116.

